# A parametrized computational framework for description and design of genetic circuits of morphogenesis based on contact-dependent signaling and changes in cell-cell adhesion

**DOI:** 10.1101/784496

**Authors:** Calvin Lam, Sajeev Saluja, George Courcoubetis, Josquin Courte, Dottie Yu, Christian Chung, Leonardo Morsut

## Abstract

Synthetic development is a nascent field of research that uses the tools of synthetic biology to design genetic programs directing cellular patterning and morphogenesis in higher eukaryotic cells, such as mammalian cells. One specific example of such synthetic genetic programs was based on cell-cell contact-dependent signaling using synthetic Notch pathways, and was shown to drive formation of multilayered spheroids by modulating cell-cell adhesion via differential expression of cadherin-family proteins. The design method for these genetic programs relied on trial and error, which limited the number of possible circuits and parameter ranges that could be explored. Here we build a parametrized computational framework that, given a cellcell communication network driving changes in cell adhesion and initial conditions as inputs, predicts developmental trajectories. We first built a general computational framework where contact-dependent cell-cell signaling networks and changes in cell-cell adhesion could be designed in a modular fashion. We then use a set of available *in vitro* results (that we call the “training set” in analogy to similar pipelines in the machine learning field) to parametrize the computational model with values for adhesion and signaling. We then show that this parametrized model can qualitatively predict experimental results from a “testing set” of available *in vitro* data that varied the genetic network in terms of adhesion combinations, initial number of cells and even changes to the network architecture. Finally, this parametrized model is used to recommend novel network implementation for the formation of a 4-layered structure that has not been reported previously. The framework that we develop here could function as a testing ground to identify the reachable space of morphologies that can be obtained by controlling contact-dependent cell-cell communications and adhesion. Additionally, we discuss how the model could be expanded to include other forms of communication or effectors for the computational design of the next generation of synthetic developmental trajectories.

## Introduction

Multicellular mammalian systems display a remarkable capacity for self-organization into a myriad of different shapes and forms, from branching lung epithelia, to elongating tail bud mesenchyme. These phenomena of self-organization in mammalian morphogenetic systems are not yet completely understood, and are being investigated intensively, for example via analysis, perturbation and modeling of model organisms^1–9^. From these studies, several factors that seem to be important for self-organization are being recognized, among them chemical, epigenetic, bioelectrical, morphogens, mechanical, cell-cell communication signals^10–12^. The identification and characterization of these factors will ultimately allow implementation in simplified “toy models” (i.e. minimal models akin to the pendulum for physics^13^) of synthetic development both *in silico* and *in vitro*^13–20^.

One paradigmatic example of morphogenetic systems is mechanochemical systems, which are composed of cell-cell signaling and changes in cellular or tissue mechanics. These systems have been shown to be at play in a number of natural systems ^21,22^, and computational models have been developed to capture both signaling and mechanical changes in integrated models ^23^.

Among mechanical effectors, cell-cell adhesion has been recognized as an important effector of multicellular morphogenesis. In its simplest form, differential adhesion between cells can favor cell rearrangements that bring cells with high cell-cell adhesion closer together. This “differential adhesion” hypothesis has been studied in cellular systems where adhesion levels were changed via constitutive adhesion protein overexpression^24^; and also via computational systems where adhesion levels can be decided by the user (including CompuCell3D^25^).

Contact-dependent cell-cell communication networks have also been recognized as powerful sources for multicellular patterning *in vivo*^26–29^, and computational models have been developed and used to show the patterning potential of contact-dependent networks^26,30,31^. Recently, synthetic variants of cell-cell contact-dependent signaling have been developed named synthetic Notch or synNotch. SynNotch is based on the native Notch receptor which relies on contact-dependent, or juxtacrine, signaling^32^. SynNotch has been engineered such that it can respond to synthetic ligands, such as green fluorescence protein (GFP), with user-defined outputs. If a cell expressing GFP-ligand (sender cell) is in contact with a cell expressing anti-GFP receptor (receiver cell), then the receiver cell will start to produce user-defined target genes. SynNotch pathways allow the user to define the input and the output of communication channels that are orthogonal to endogenous signaling. This synthetic cell-cell contact signaling system alone has been used to drive patterning in 2D in epithelial cells^33^.

A first example of a synthetic artificial genetic networks for mechanochemical-driven morphogenesis was recently generated by combining cell-cell adhesion and cell-cell contact mediated communication^19^. In this system, artificial genetic networks intertwine two morphogenetic factors: cell-cell contact-dependent signaling and concomitant changes in cellcell adhesion. Changes of cell-cell adhesion are initiated via synNotch cell-cell communication pathways, leading to dynamic and localized changes in cell-cell adhesion strengths. With these networks, mouse fibroblast cells can be designed such that they follow user-defined morphogenetic trajectories to generate multilayered spheroids starting from random mixture of 1 or 2 genetically different cell types. This work demonstrates that simple networks of cell-cell communications with changes in cell adhesion can drive developmental trajectories that have features commonly found in developing morphogenetic systems.

One limitation of synthetic development studies lies in the design phase which relies heavily on lengthy trial-and-error iterations. As the range of parameters and designs that can be tested is limited to scientists’ best guesses, a systematic analysis of design space is not possible and interesting solutions may be overlooked. In other areas of synthetic biology this initial phase of intuitive design has been followed by a phase of computational systems development. These computational systems are used initially for the description and then the design of the systems in a way that can lead to implementation^34–36^. One remarkable example is the decade-long development of Cello, a computational framework for design and implementation of bacterial unicellular gene regulatory networks^37^. In Cello, users can give design specifications *in silico* of combinatorial logic, and the computational system converts them into complete DNA sequences encoding transcriptional logic circuits that can be executed in bacterial cells. Similar efforts in mammalian cells are happening with a lag, beginning with intracellular circuits^38^. Computational efforts for helping the design of user-defined structures in multicellular systems have been initiated, for example the design of structure obtained by sculpting of muscle and epithelial cells in “xenobots”^39,40^, or the design of culture and initial conditions for aggregates of human stem cells to control their organization^41^. Until recently, no example of a computational system for description at the level of genetic circuits of morphogenesis was available for assisting with design. The *in vitro* system in Toda seems perfectly placed to provide a case study as it has all the features of a minimal “toy model” of synthetic circuit-guided morphogenesis (and it has attracted efforts from other groups as well^42,43^): it has logically minimal circuits, shows the genotype-to-phenotype relationships, and has explored the phenotypic consequences of changes in either cell-cell adhesion and/or network topologies.

Here we describe how we have built a parametrized computational framework for the description and design of genetic circuits of morphogenesis based on contact-dependent signaling and changes in cell-cell adhesion. This computational system can take, as input, an artificial genetic circuit of cell-cell communication, changes in cell adhesion (a “synthetic genome”) and initial conditions (e.g. number of cells of each type), and produce, as output, the developmental trajectory (Fig. 1A-B). The computational system is based on a stochastic cellular Potts environment implemented in CompuCell3D, which allows the simulation of adhesion-based rearrangements, cell movement and cell division^25^. We overlaid custom code to model cell-cell-contact dependent signaling by defining cell types and encoding information about cell-cell contact. We then we split the available *in vitro* dataset into a “training set” and a “testing set” (Fig. 1C), following a framework for model identification common in machine learning. We use the “training set” to identify parameters for the *in silico* model. The synthetic genomes of the training set are chosen such that they would contain all the basic primitives of signaling and adhesion interactions that are found in the *in vitro* implementations: adhesion strengths, cell movement and proliferation, and signaling.

**Fig. 1.**
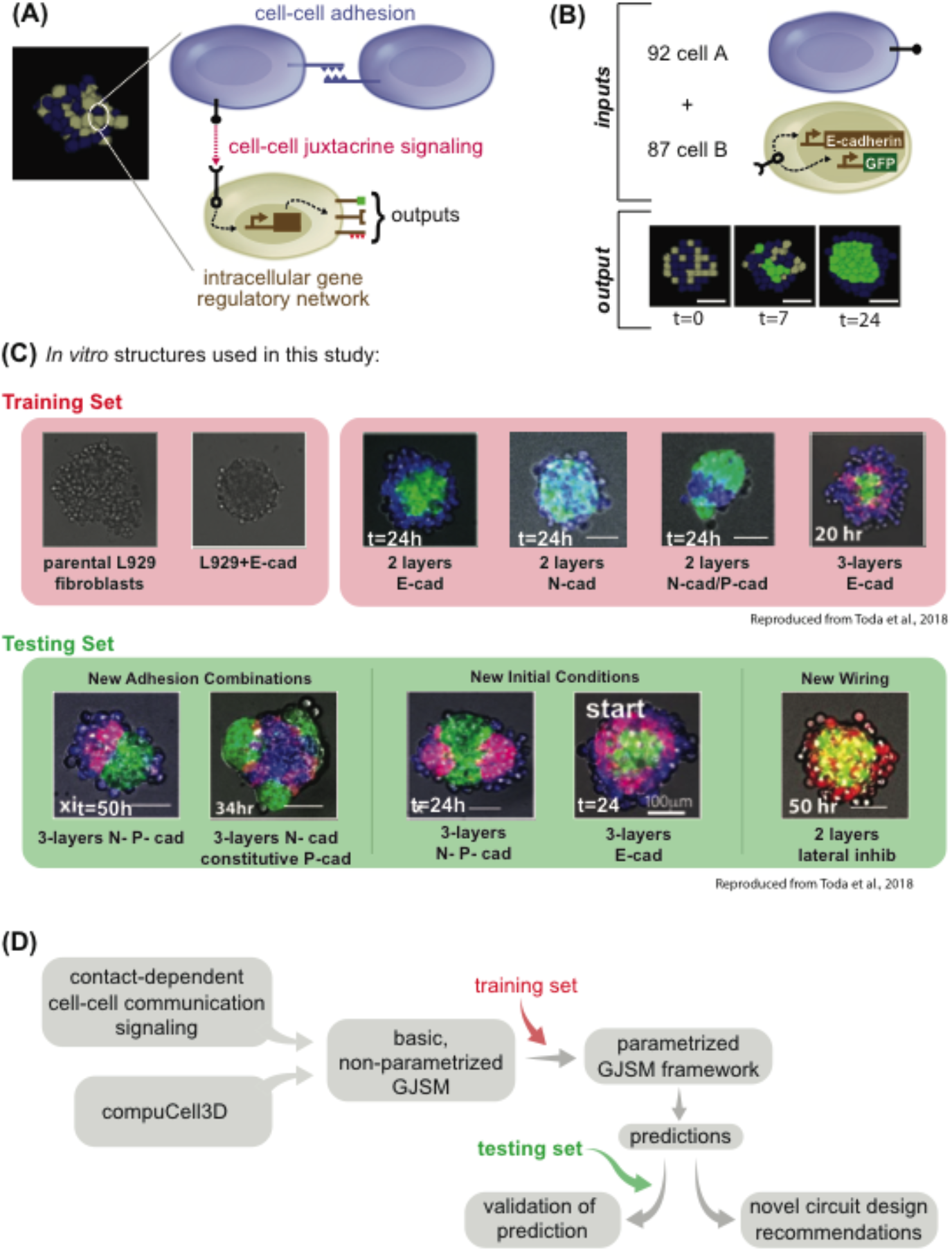
The model is developed and tested via a training set/testing set approach to parametrize and test generalization capacity. **(A)** Example of a snapshot of computational model output on the left; highlighted on the right is the schematic of the logic that drives the cell dynamics: cell-cell adhesion, and cell-cell juxtacrine signaling combined together. Cells can communicate to one another if, as in this case of blue type and gray type, have the cognate signal and receptor. Outputs of the communication can be a change in fate which can signify: new ligands production or changes in cell adhesion. **(B)** We define input to our computational model as: the initial conditions of how many cells of the different types there are, and the “genotype” of cells of the different types; the output is a simulation of the developmental trajectory. **(C)** Snapshot of the in vitro structures used in this study to build the computational model. The larger black box encompasses all the structure; the red box highlights the structures used for the training set; the green box the structures for the testing set. For each structure, a microscopy image of L929 cells engineered with different networks is shown, as described later in the text, and imaged at the indicated timepoint. 2-layer indicates a network of signaling where A➔B, and 3-layer one where A➔B➔A (see later for more details). Adhesion molecules are as follows: Ecad= E-cadherin; Ncad = N-cadherin; Pcad = P-cadherin. Scale bar is 17.5pixels *in silico*, 100um *in vitro*. **(D)** Conceptual diagram of the flow we followed in the entire paper: each box represents a conceptual item that we used and combine logically. First, we generate a basic GJSM by combining CompuCell3D with custom code for cell-cell contact dependent signaling and change in cell types; then we use the training set to identify parameters capable of recapitulating in vitro training set behaviors; finally we use that model to make prediction, which are tested by comparing them with the testing set, or used to make recommendations for novel synthetic genomes. Microscope images with gray background are reproduced from^19^, reprinted with permission from AAAS.

Subsequently, we use the testing set to test if parametrization is able to capture features of the systems outside the ones used for parametrization. Finally we identify a synthetic genome *in silico* that can generate novel 4-layer structures (not present in the *in vitro* dataset) and provide a recommendation for the synthetic genome that could implement it *in vitro* with a combination of available parts (Fig. 1D). Figures 2–4 describe the model parametrization based on *in vitro* data. Figures 5–8 show examples of the capacity of the parametrized computational system to quantitatively reproduce emergent properties not used in parameterization (Fig.5) and qualitatively predict developmental trajectories from the testing set that have different adhesion (Fig. 6), different initial number of cells (Fig. 7), or changes to the genetic network (Fig. 8). Figure 9 shows the recommendation for the synthetic genome for the 4-layered structure *in vitro* with a combination of available parts.

**Fig. 2.**
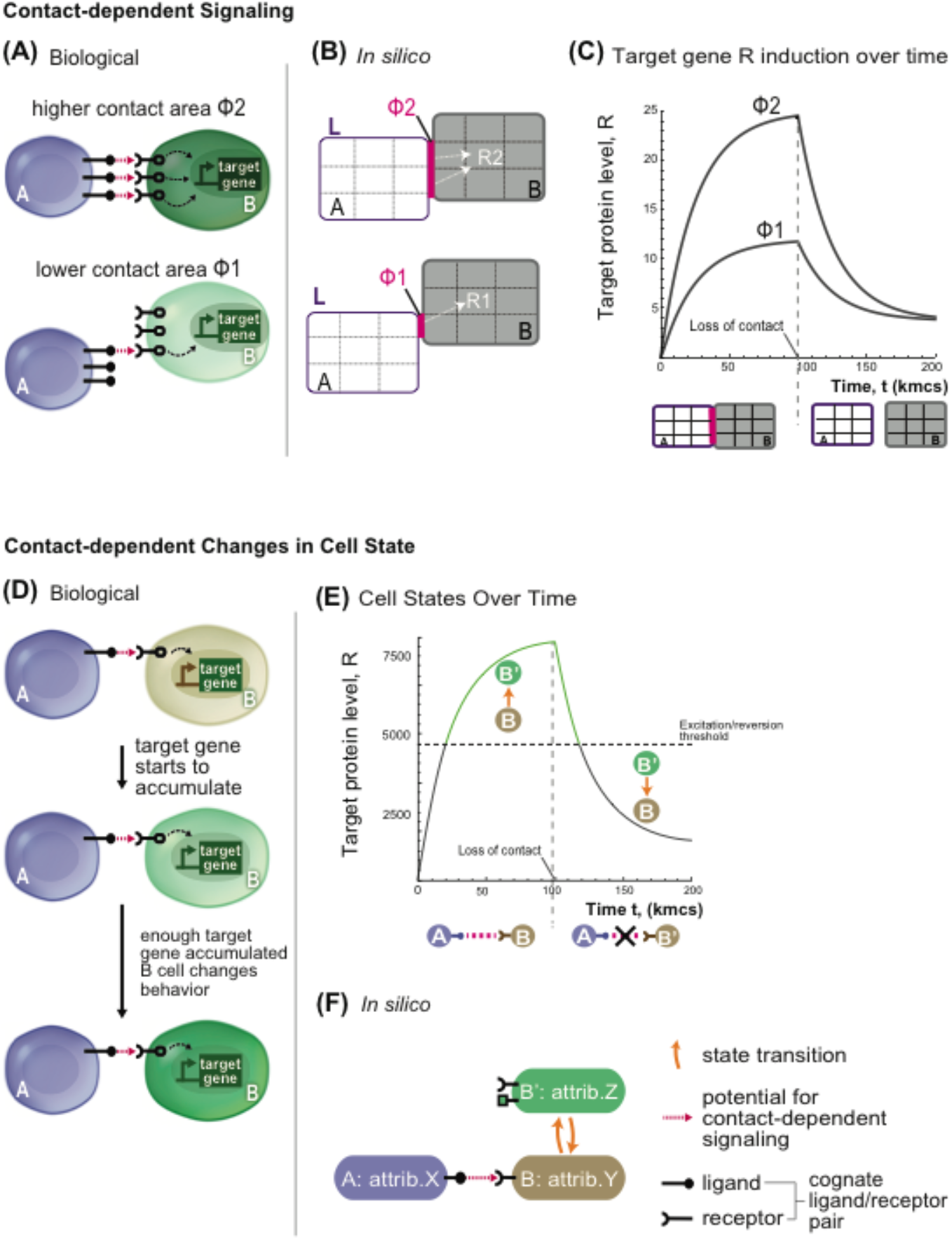
Concepts underlying the computational model (See Methods for details and generalized model). Schematic description of computational model for contact-dependent cell state changes in cell aggregates. **(A)** Representation of biological communication between cell pairs A and B. A cells express ligand (purple circle + stick) and B cell express receptor (black). With contact (pink arrow), B cells receive signal (dashed arrow) that triggers expression of the target gene (green). In the cells below, the amount of contact surface is lower, hence the signaling (dashed arrow) towards the target gene is less intense. **(B)** The *<u>in silico</u>* model shows a simplified representation of the cell-cell signaling process with param eters: ligand amount (L, purple), surface area of contact (Φ, pink), and net signal (white arrow to target gene R). Cells (A) are the sender in the communication, cells B are the receiver. In this schematic, *in silico* cells are objects of 9 pixels. The cell pair at the top has a higher level of signaling compared to the upper pair due to a larger surface area of contact (Φ2>Φ1, 2 pixels compared to 1). **(C)** Time evolution of target gene level in the receiving cell B; cells A and B are placed in contact at time zero and kept in the same configuration for 100,000 steps of simulation, and then moved far apart to stop signaling. Target gene levels is followed over time for two different values of shared surface area Φ, with Φ2>Φ1, with all other parameters kept identical. **(D-F)** Model representation of cell behavior state change. **(D)** Representation of biological “effector gene” activation: a sender cell A (blue) activates a receiver cell B (gray) to induce a target gene (green) that encodes for an effector protein. Over time, cell B accumulates target genes products, and at a certain threshold the effector gene product causes cell behavior to change (e.g. stronger adhesion to neighbors) such that cell transition from type B to B’. **(E)** The graph shows the progression of target gene level over time for a B cell that is initially in contact with an A cell and is then isolated at 100,000 steps. Threshold for the excited state is shown as dotted horizontal line. At the start, the B cell is in the basal state (black solid line), but when the target gene level passes the excited state threshold, B cell becomes a B’ cell. The B’ cell remains in the active state (green solid line) until target gene levels drop below the activation threshold and reverts to B (line goes back to solid black). **(F)** *In silico* representation of the state transition and communication relationship between cells A, B, and B’. Orange curved arrows indicate state transitions. Matching ligand/receptor pairs indicate a communication channel from A to B that promotes the state change of B to B’.

**Fig. 3.**
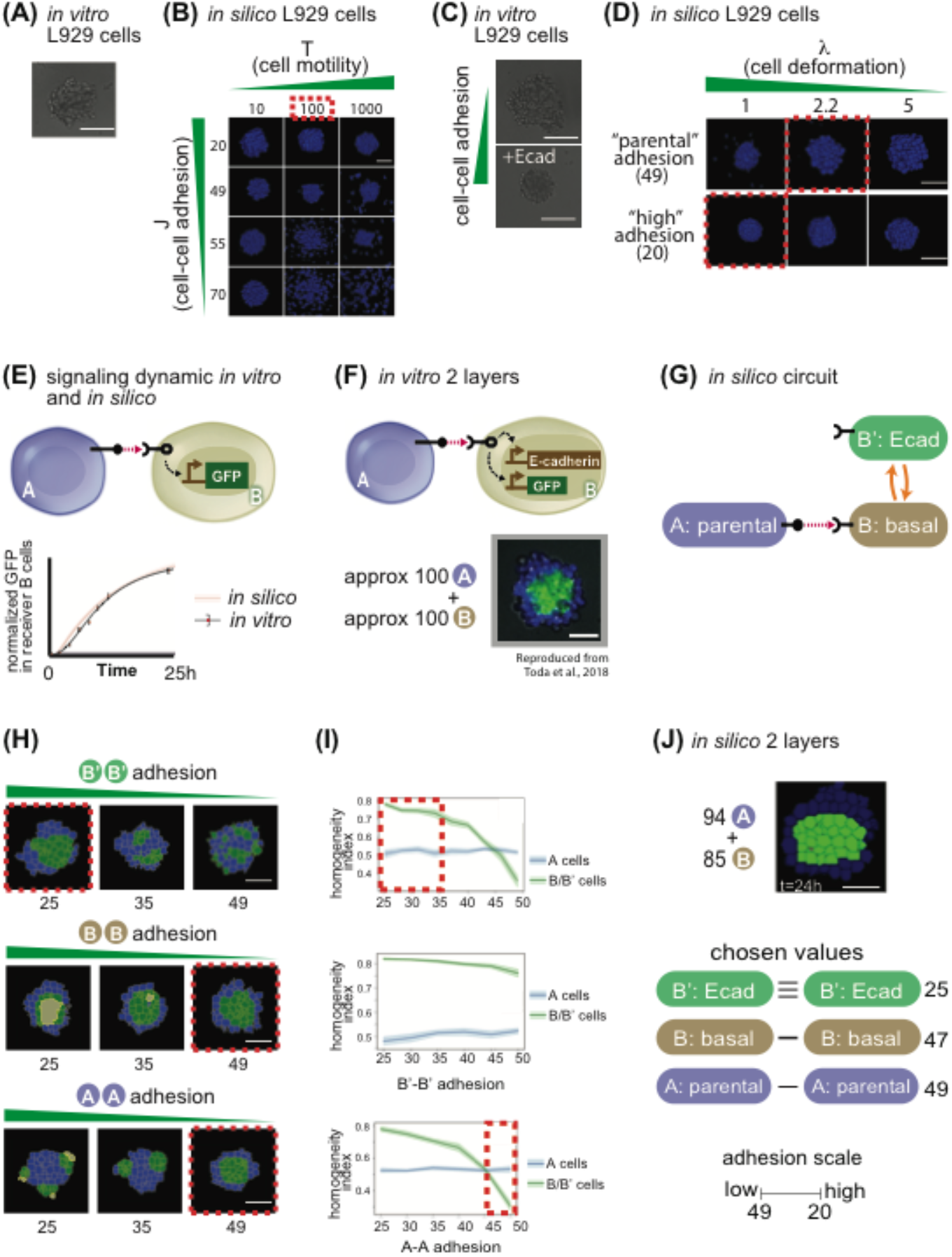
Model parametrization of signaling and adhesion in A➔B networks. **(A)** In vitro picture of 100 parental L929 cells grown for 24h in a non-adhesive U-bottom well of a 96-well plate. **(B)** In silico pictures of 100 identical cells grown in a non-adhesive virtual medium for 24h; as indicated, cells in different snapshots have different parameters for cell movement (x axis) and cell-cell adhesion (y axis). **(C)***In vitro* picture of 100 L929 cells grown for 24h in a nonadhesive U-bottom well of a 96-well plate, either parental (upper picture) or genetically engineered to overexpress E-cadherin (lower picture). **(D)***In silico* pictures of 100 identical cells grown in a nonadhesive virtual medium for 24h; as indicated, cells in different snapshots have different parameters for cell deformation (x axis) and cell-cell adhesion (y axis). **(E)** Diagram of sender A cells, and receiver B cells that induce GFP downstream of activation of a contact-dependent receptor (left). Right, graph of target gene expression over time for B cells for either in silico simulations (red line + shadow standard deviation) or in vitro experiments (black line). **(F)** Diagram of sender A cells, and receiver B cells that induce GFP and E-cadherin downstream of activation of a contact-dependent rec eptor (left). On the right is a result of an in vitro experiment after 24h of cultivating approximately 100 A cells with 100 B cells. **(G)** Depiction of transition network between cell states that is implemented in Fig. F. **(H)**, starting from 100 A and 100B cells where the B’-B’ adhesions where changed (first line), or the A-A (second line) or the B-B (third line). Red dotted lines indicate structure that most closely resemble the in vitro implementation (F). **(I)** Sorting index quantification of a 24h timepoint of cells A (blue line) or B (green line), for a range of B’-B’ adhesion (first line), A-A adhesions (second line) and B-B adhesions (third line). Red dotted lines represent ranges of behavior that recapitulate in vitro observations. **(J)***In silico* output of input 94 A cells and 85 B cells with the same network as in H, and with the values for adhesion indicated on the right. Scale bar is 17.5pixels *in silico*, 100um *in vitro*. Microscope images with gray background are reproduced from^19^, reprinted with permission from AAAS.

**Fig. 4.**
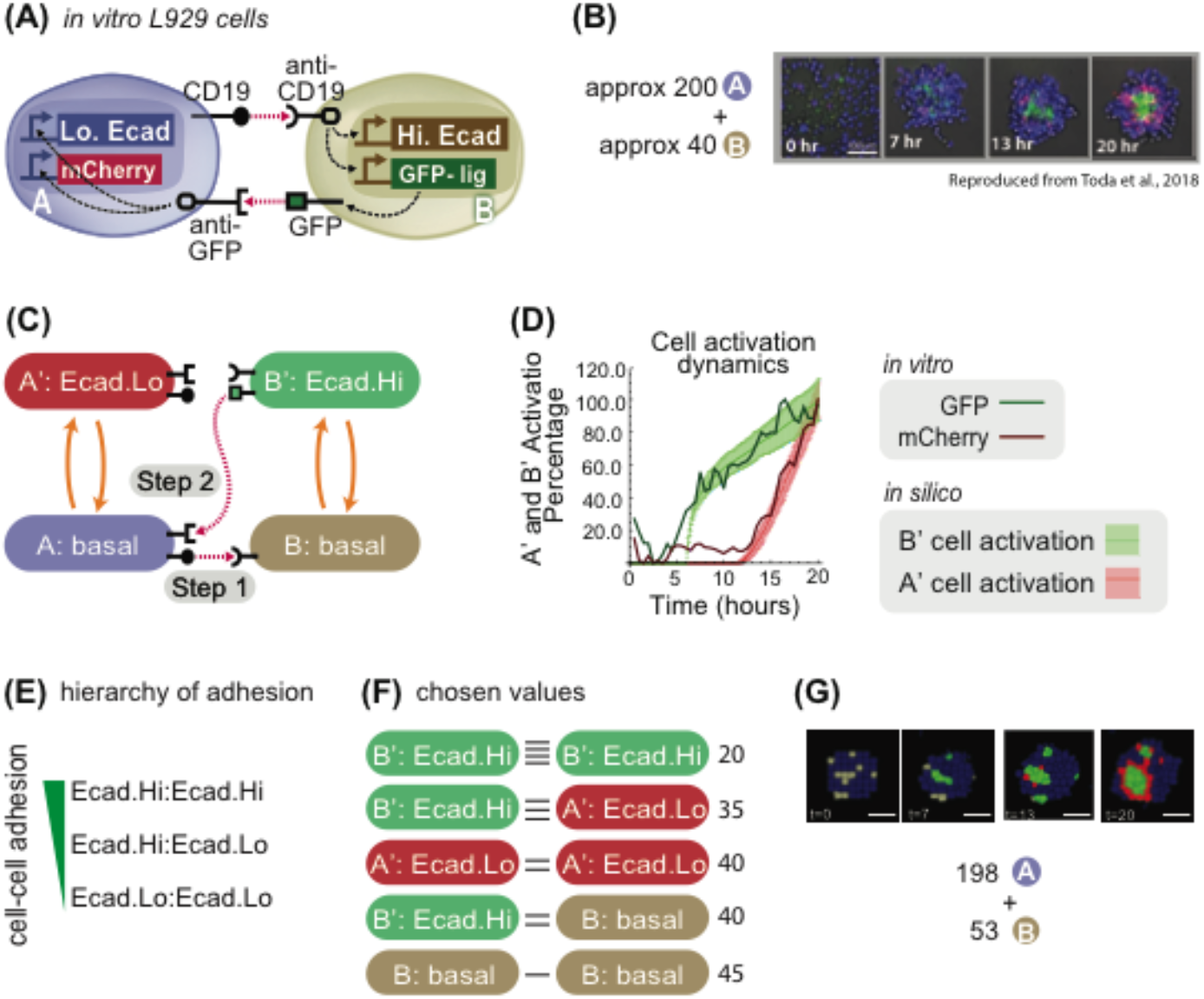
Parametrization of A➔B➔A network parameters and E-cadherin levels. **(A)** Diagram 2 cells A (blue) and B (gray) and their signaling with the receptor and promoter representation: cell A has a CD19 ligand (rounded ligand on stick on cell membrane of cell A), which activates (pink arrow) rounded receptor in cell B, which in response activates Hi.Ecad (white rectangle) and GFP-ligand (green rectangle); the GFP-lig (green square on a stick on the membrane) is then produced (gray arrow) on the membrane, which activates the square receptor on cell A that activates intracellularly (white arrows) Lo.Ecad (white rectangle) and mCherry (red rectangle). **(B)** Experimental results of mixing approximately 200 A cells and 40 B cells; A cells are blue, B cells are gray, B’ cells are green and A’ cells are red. **(C)** State-machine diagram of the network, with the adhesion components. B cell has basal adhesion, and rounded receptor; when it is activated by a neighboring rounded receptor it activates to a B’ state where it starts to be adhesive with Ecad.Hi parameter, and a square ligand; A cell, blue, has basal adhesion, expresses rounded ligand, and a square receptor; the activation of the square receptor activates A cell to A’, which is red and acquire adhesion of Ecad.Lo parameter. **(D)** Graph depicting cell type activation index over time, for activation from A➔A’ in vitro (dark red line) and in silico (light red line and shadowed standard deviation) and from B➔B’ in vitro (dark green line) and in silico (light green line and shadowed standard deviation). We present mean±s.d. for the *in silico* results (dotted lines with standard deviations in the graph). (n=30 simulations *in silico*, n=1 for *in vitro*). **(E)** Hierarchy of adhesion between cells with different levels of E-cadherin adhesion; notation adhesion.level1 : adhesion.level2 indicates adhesion preference between cells that express adhesion level 1 and cell that express adhesion level 2. **(F)** Diagram of the cell-cell adhesion strengths for pair-wise cells of different types; horizontal black lines denote cell-cell adhesion, and more horizontal line denote stronger adhesion. **(G)** Input (198A + 53B cells) and output (developmental trajectory, with snapshots at the indicated time frames) of the system with genome as in (C). Scale bar is 17.5pixels *in silico*, 100um *in vitro*. Microscope images with gray background are reproduced from ^19^, reprinted with permission from AAAS.

**Fig. 5.**
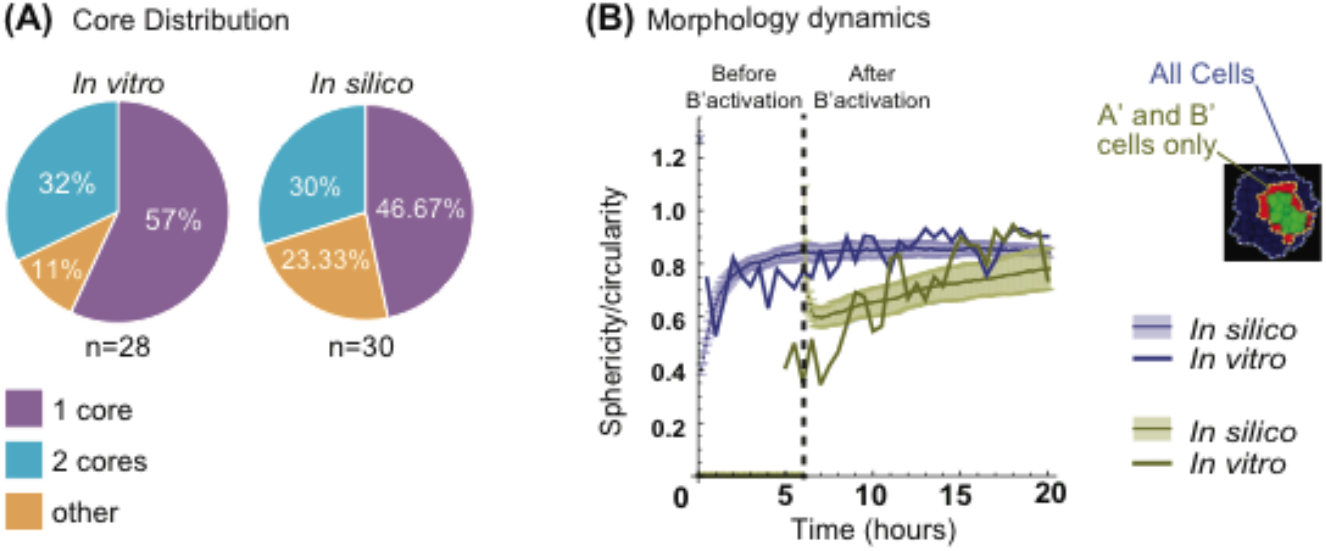
Emergent properties in vitro are captured by the parametrized system in silico for the central symmetric three layers network. **(A)** Quantification of the number of cores formed over repeated simulations (n=30 simulations, n=28 for *in vitro*). *In vitro* results data are from^19^. See Methods, “Quantification and Statistical Analyses; Simulation Quantifications”, for more details on cores definitions *in silico*. **(B)** Quantification of sphericity/circularity indexes over the time development of synthetic and *in vitro* systems. (See methods, Simulation quantifications for *in silico*, and Video Analysis for *in vitro* details on the indexes). In blue, all the cells are considered; in green only the activated (A’) and (B’) cells. Solid line is from *in vitro* measures; solid lines with shaded contours are from *in silico* measurements and represent mean and standard deviation interval respectively. Vertical dashed line indicates time of (B’) cells activation (n=30 simulations, n=1 for *in vitro*).

**Fig. 6.**
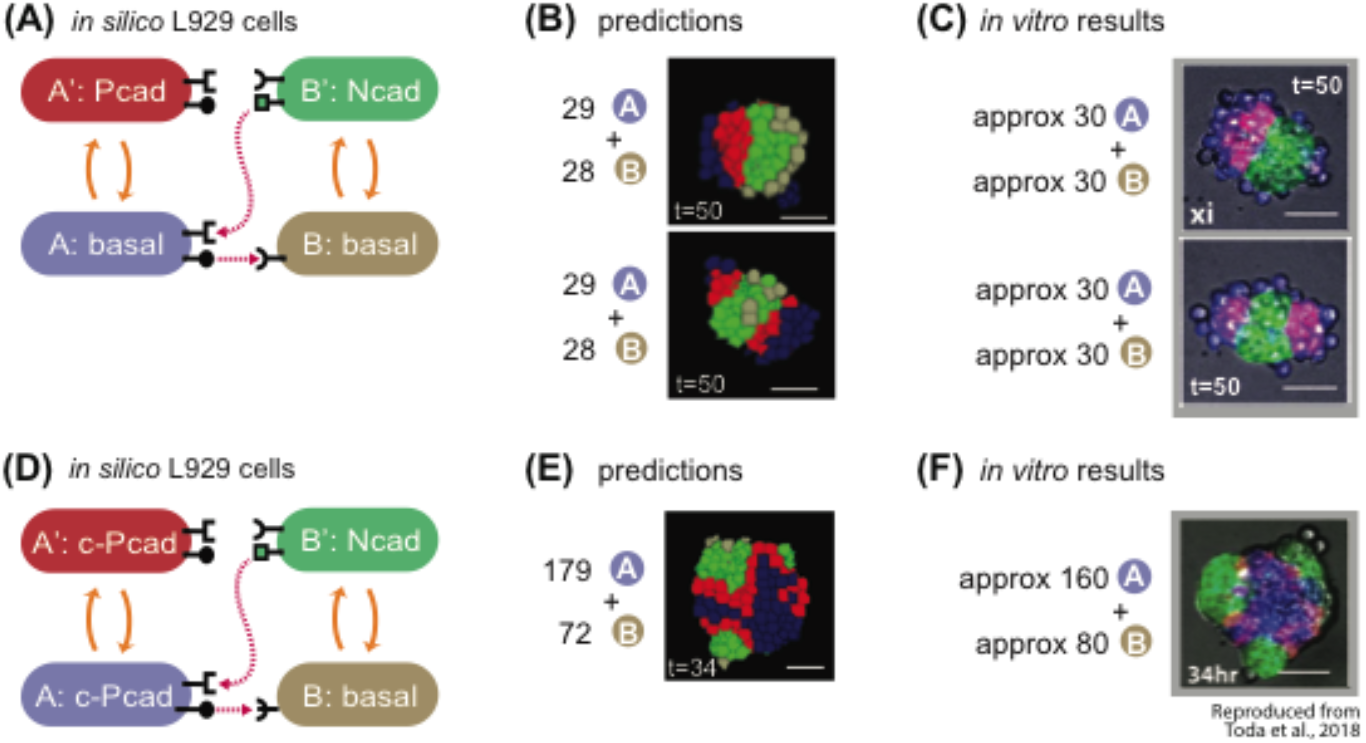
The model correctly predicts behavior of in vitro experiments with different cadherin target genes. **(A)** State-machine diagram of the network, with the adhesion components. B cell has basal adhesion, and rounded receptor; when it is activated by a neighboring rounded receptor it activates to a B’ state that is adhesive with Ncad parameter, produces a square ligand; A cell, blue, has basal adhesion, expresses rounded ligand, and a square receptor; the activation of the square receptor activates A cell to A’, which is red and has adhesion of Pcad parameter. **(B)** Representative results of 2 classes of resulting structures at t=50h of simulated time of approximately 30A + 30B cells with the genotype as in (A): one class (approx. 10% of the simulations) gives 2 red poles, and the other class (approx. 90% of the simulations) gives 1 red pole. See Supplem. Fig. S3 for quantification of homogeneity index over 10 runs of the simulation with the same parameters and initial conditions. **(C)** In vitro results at t=50h of the 2 category of structures that are obtained with the genotype of cells in A, and with mixing approximately 30A cells and 30B cells. **(D)** State-machine diagram of the network, with the adhesion components. B cell has basal adhesion, and rounded receptor; when it is activated by a neighboring rounded receptor it activates to a B’ state that is adhesive with Ncad parameter, produces a square ligand; A cell, blue, has constitutive Pcad adhesion, expresses rounded ligand, and a square receptor; the activation of the square receptor activates A cell to A’, which is red and has continues to have adhesion of constitutive Pcad parameter. **(E)** Representative results of resulting structures at t=34h of simulated time of approximately 179A + 92B cells with the genotype as in (D): a core of blue inactivated A cells, with multiple poles of green activated B’ green cells, and in the between red activated A’ cells. See Supplem. Fig. S3 for quantification of homogeneity index over 10 runs of the simulation with the same parameters and initial conditions. **(F)** in vitro results at t=34h of a representative structure that is obtained with the genotype of cells in (D), and with mixing approximately 160A cells and 80B cells. Scale bar is 17.5pixels =100um. Microscope images with gray background are reproduced from^19^, reprinted with permission from AAAS.

**Fig. 7.**
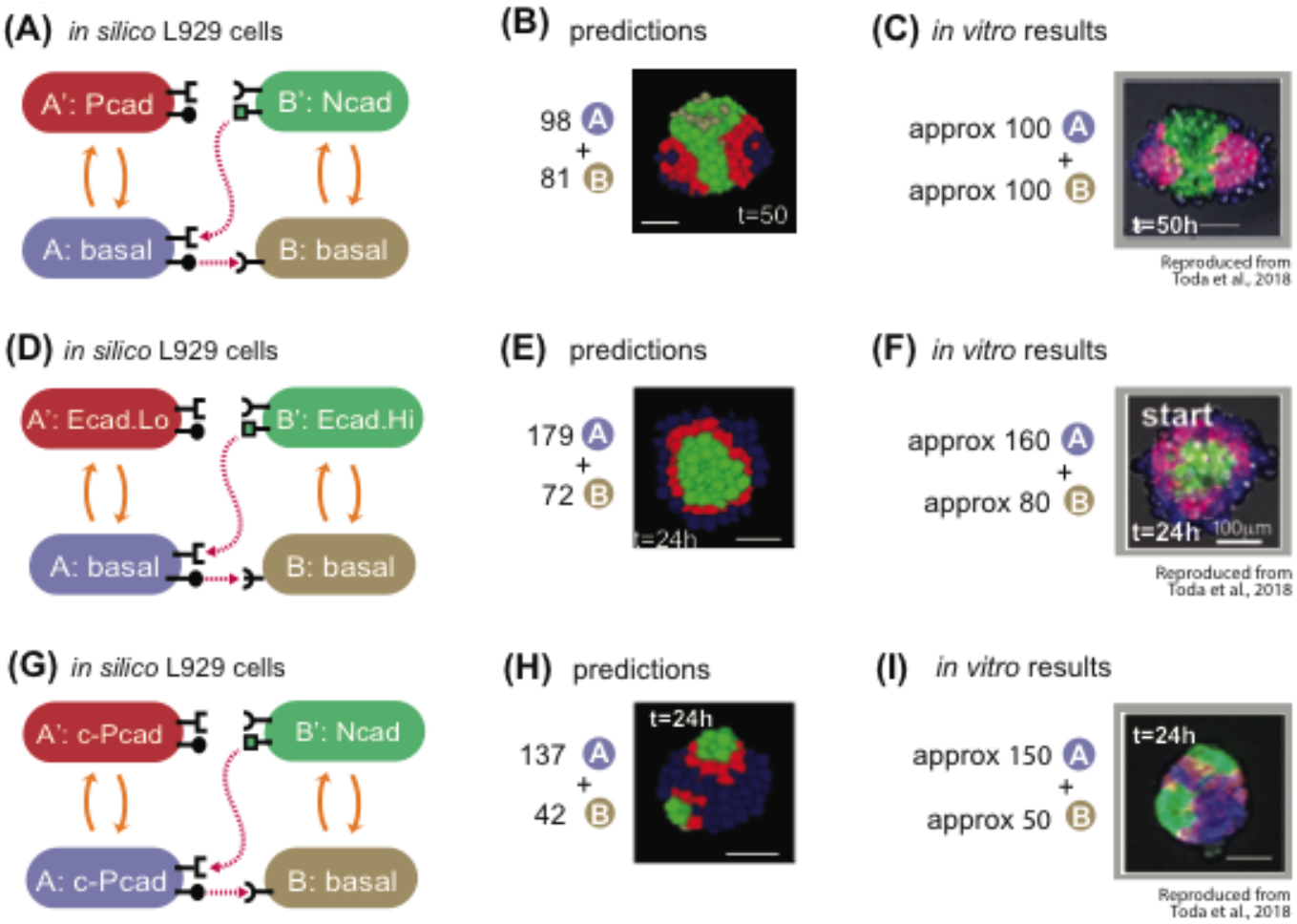
The model correctly predicts behavior of *in vitro* experiments with different number of cells. **(A)** State-machine diagram of the network, with the adhesion components. B cell has basal adhesion, and rounded receptor; when it is activated by a neighboring rounded receptor it activates to a B’ state that is adhesive with Ncad parameter, produces a square ligand; A cell, blue, has basal adhesion, expresses rounded ligand, and a square receptor; the activation of the square receptor activates A cell to A’, which is red and has adhesion of Pcad parameter. **(B)** Representative results of resulting structures at t=50h of simulated tim e of 98A + 81B cells with the genotype as in (7A): a core of green activated B’ cells, with two poles of red activated A’ cells and external to that blue inactivated A cells. See Supplem. Fig. S4 for quantification of homogeneity index over 10 runs of the simulation with the same parameters and initial conditions. **(C)***In vitro* results at t=50h of a representative structure that is obtained with the genotype of cells in (7A), and with mixing approximately 100A cells and 100B cells. Scale bar is 17.5pixels =100um. **(D)** State-machine diagram of the network, with the adhesion components. B cell has basal adhesion, and rounded receptor; when it is activated by a neighboring rounded receptor it activates to a B’ state that is adhesive with Ecad.Hi parameter, produces a square ligand; A cell, blue, has basal adhesion, expresses rounded ligand, and a square receptor; the activation of the square receptor activates A cell to A’, which is red and has adhesion of Ecad.Lo parameter. **(E)** Representative result of resulting structures at t=24h of simulated time of approx. 179A + 72B cells with the genotype as in (7D): a core of green activated B’ cells, with a subsequent layer of red A’ activated cells, surrounded by a layer of inactivated blue A cells. See Supplem. Fig. S4 for quantification of homogeneity index over 10 runs of the simulation with the same parameters and initial conditions. **(F)***In vitro* results at t=24h of a representative structure that is obtained with the genotype of cells in (7D), and with mixing approximately 160A cells and 80B cells. Scale bar is 17.5pixels =100um. **(G)** State-machine diagram of the network, with the adhesion components. B cell has basal adhesion, and rounded receptor; when it is activated by a neighboring rounded receptor it activates to a B’ state that is adhesive with Ncad parameter, produces a square ligand; A cell, blue, has constitutive Pcad adhesion, expresses rounded ligand, and a square receptor; the activation of the square receptor activates A cell to A’, which is red and has continues to have adhesion of constitutive Pcad parameter. **(H)** Representative results of resulting structures at t=24h of simulated time of 137A + 42B cells with the genotype as in (G): a core of blue inactivated A cells, with multiple poles of green activated B’ green cells, and in the between red activated A’ cells. See Supplem. Fig. S4 for quantification of homogeneity index over 10 runs of the simulation with the same parameters and initial conditions. **(I)***In vitro* results at t=24h of a representative structure that is obtained with the genotype of cells in (G), and with mixing approximately 150A cells and 50B cells. Scale bar is 17.5pixels =100um. Microscope images with gray background are reproduced from ^19^, reprinted with permission from AAAS.

**Fig. 8.**
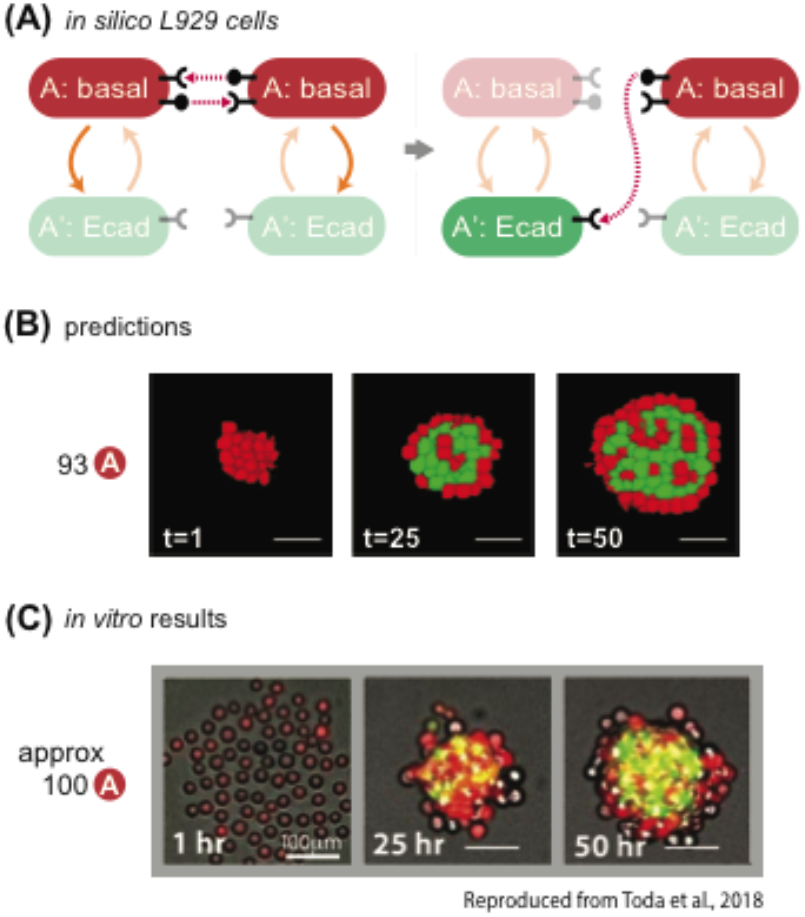
The model co rrectly predicts behavior of in vitro experiments with different network wiring. **(A)** State-machine diagram of the network, with the adhesion components. Cells of type A are red and have basal adhesion and express both rounded ligand and receptor; when the receptor is activated enough, it can activate A cells to A’ cells, which lose rounded signal expression, gain color green and expression of Ecadherin. **(B)** Representative results of resulting structures at the indicated timestamp of developmental trajectory starting from 93 cells of type A with the genotype as in (8A); some of the red cells turn green and gather at the center of the core; at t=50, the external layer is of inactivated A cells, and the internal core is a mixture of mainly green cells with interspersed a minority of red cells. See Supplem. Fig. S5 for quantification of homogeneity index over 10 runs of the simulation with the same parameters and initial conditions. (**C**) *In vitro* results at the indicated time points of a representative structure that is obtained with the genotype of cells in (8A), starting with 100 A cells. Scale bar is 17.5pixels =100um. Microscope images are reproduced from ^19^, reprinted with permission from AAAS.

**Fig. 9.**
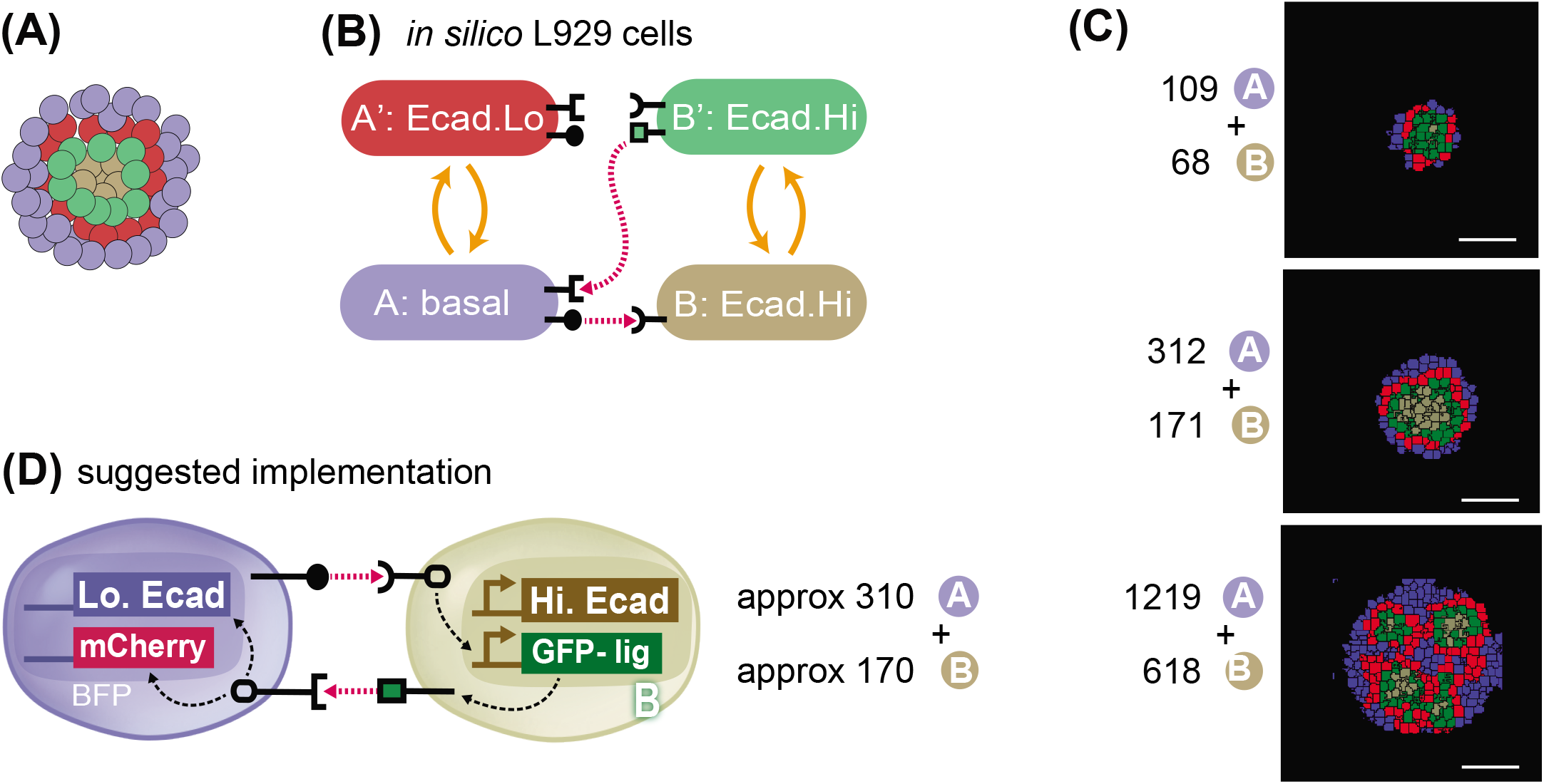
*In silico* recommendation for obtaining a 4-layered structure. **(A)** Goal target pattern: 4 layers with gray-green-red-blue cells centrally symmetric. **(B)** State-machine diagram of the network, with the adhesion components. B cell has adhesion with Ecad.Hi parameter, and rounded receptor; when it is activated by a neighboring rounded receptor it activates to a B’ state that is adhesive with the same Ecad.Hi parameter, and produces a square ligand; A cell, blue, has basal adhesion, expresses rounded ligand, and a square receptor; the activation of the square receptor activates A cell to A’, which is red and has adhesion of Ecad.Lo parameter. **(C)** Representative results of resulting structures at t=24h of simulated time of cells with the genotype as in (9A) and with the ratio and numbers as indicated. All structures generate inactive A cells, activated A’ cels, inactive B cells and activated B’ cells in different ratios and in different geometrical arrangements. See Supplem. Fig. S6 for quantification of homogeneity index over 10 runs of the simulation with the same parameters and initial conditions. **(D)** Recommended *in vitro* implementation. Scale bar is 17.5pixels *in silico*, equivalent to approx. 100um *in vitro*.

This example of computational pipeline is among the first examples of computational systems used for description and design of genetic circuits for synthetic morphogenesis based on synNotch and changes in adhesion. We think that it will provide a first necessary step towards the development of complete pipelines with computational design and *in vitro* implementation with this specific toolkit, as well as provide inspiration for extension of this framework to other methods of communication and morphogenetic effectors (e.g. soluble ligand based, bioelectricity, cytoskeletal, etc.).

### Framework for modeling of cell-cell contact signaling and cell state changes in CompuCell3D (complete details in Methods)

We first built a computational framework that could simulate development of multicellular spheroids informed by a “synthetic genome” that controls their cell-cell communication, adhesion, movement. The CompuCell3D platform can simulate adhesion, movement, division and presence of different cell “types”, we added new code for contact-dependent signaling mediated change of cell type.

Briefly, in CompuCell3D^25^, each cell *σ*(*i*) is defined as comprising a user-defined number of atomic pixels, whose position in space can change over time with a stochastic algorithm. The probability that a pixel will move is calculated by CompuCell3D for each attempt of movement at each time interval (known as Monte Carlo step (mcs)) with the following equation:

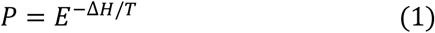

Where ΔH is the difference in “entropy energy” between the configurations before the change versus after the change, and T is a “temperature” parameter of the cell that is attempting the change which can be interpreted as cell motility. Governed by this equation, a system will dynamically change configuration over time towards configurations that minimize entropy energy (H). Entropy energy for a given a configuration of cells is calculated based on cell-cell contact and cell shape as follows:

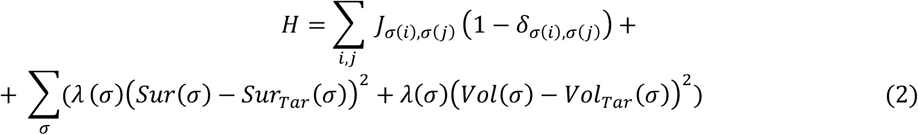

The first term of this equation deals with how cell-cell contacts contribute to H. Whether two cells are in contact is encoded in a Kronecker delta *δ*_*σ*(*i*),*σ*(*j*)_ that is 0 if cells i and j are neighbors, and 1 otherwise and can be thought of as an adjacency matrix. If two cells are in contact, they will contribute proportionally to H the number of pixels that are in contact weighted by the adhesion matrix J. The adhesion matrix *J*_*σ*(*i*),*σ*(*j*)_ defines an adhesion weight between cell i and j. Higher adhesion values in the adhesion matrix correspond to increased entropy, less “stable”conformations, and hence a lower likelihood that neighboring cells will remain close.

Consequently, higher adhesion values in the model correspond to a lower likelihood of cells sticking together and can therefore be interpreted as corresponding to lower adhesion strength. The second line of equation (2) deals with how cell shape contributes to H. Cell shape contributions are defined as the sum of all deviations of current volume of each cell from target volume (VolTar) and surface (SurTar), weighted with a “deformability” parameter λ. As cells deform and stray further from their target volume or surface, entropy energy (H) increases. The amount that the deformation is penalized is weighted via λ, with lower values of λ allowing larger deformations.

In this context, cell types can be defined as subsets of the total set of cells that share certain features (e.g. adhesion, motility, deformability, or other). Upon cell division, daughter cells inherit the features of the parent cell. One special “cell type” indication is given to the medium, such that cell-medium adhesion parameters can be defined.

In this framework, we introduced the capacity for cells to be able to influence the behavior of their neighbors, so that a contact-dependent cell-cell communication system could be implemented. The abstract features of the synNotch communications that we want to capture are: (i) signaling that is proportional to amount of shared cell-cell contact surface between sender and receiver cells that express cognate ligand/receptor pairs, (ii) signaling that is capable of affecting changes in protein production, (iii) that when protein production rises sufficiently, the receiver cell can change from a basal to an activated state, and (iv) that the activated cell can acquire new behaviors such as altered adhesiveness, or the capacity to send a new signal (Fig. 2A,D). To model these synNotch communications, we conceptually separated the modelling into two parts: signaling-dependent continuous changes in target protein production in receiver cells (Fig. 2B-C), and protein-dependent discrete changes in cell behavior (Fig. 2E-F). The two parts of the model, continuous signaling and discrete response, are highly modular and can be designed and tested independently of one another.

For modeling <u>contact-surface dependent, continuous changes in target protein production</u>, we introduce a new feature of *in silico* cells, their repertoire of ligands and receptors. Each cell *σ*(*i*) can be equipped with ligand_A through _Z, and with receptor_A through _Z. If a cell is equipped with receptor_X, it can start to accumulate target protein production points if it is in contact with a cell that is expressing the cognate ligand_X. Cell contact information is encoded in the adjacency matrix (see above) and the response is calculated via differential equations to model input-dependent response as follows:

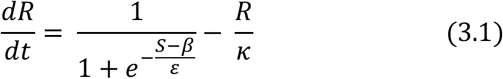

here R is the response (target protein production); S is the signal, or input coming from neighbors; β, is a constant that controls signaling delay; ε is a constant that affects the strength, steepness and overall geometry of this differential equation; k is the degradation constant. Signal (S) can depend on several factors: the number of sender cells contacting the receiver cell, the number of ligands on each sender cell, the number of receptors on the receiver cell, and the amount of contact between sender(s) and receiver. In a simplified two-cell case with sender cell A and receiver cell B, if the receptors on B are in excess, signaling depends primarily on the amount of ligand on cell A and the fraction of A’s surface contacting B. If we define L as the number of ligands on cell A’s surface and Φ as the fraction of A’s surface in contact with B’s surface, we can then define the signal (S) that cell B receives as S=Φ*L (Fig. 2B).

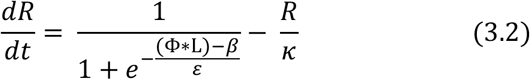

The target protein production for each cell is calculated by numerically solving (3.2) via the forward Euler method. With these definitions, we have cellular signaling that depends on the amount of contacted ligand and obtain a stronger response as the fraction of shared surface or ligand produced per unit area are increased, as in the case of synNotch signaling^33,44^ (Fig. 2C). This part of the model accounts for the continuous changes in protein production in receiver cells. For a more detailed explanation, please see the “CompuCell3D and the cellular Potts Formalism” and the “Generalized Juxtacrine Signaling Model (GJSM)” section of the Methods.

To model <u>target protein-dependent change in cell behavior</u>, we define thresholds of protein production that induce transition from a basal cell state to an activated state and vice versa. In the example above, if receiver cell B receives signal that will allow to produce enough protein to pass the activation threshold the cell will become activated, which we denote as B’ (Fig. 2E). Thresholds for state transitions from basal to active and from active to basal can be different in general, but in the current work are kept at the same value. The state machine schematic for this type of network is in Fig. 2F. The features of an activated cell’s adhesion and signaling can be different from its basal state. For example, B’ cells can be more adhesive than B cells to other B’ cells. In this way, behavioral transitions are linked to the protein production changes via a threshold, which captures the discrete change in cell behavior observed in cell biology and used in other modeling efforts^3,4,45^. Additionally, or alternatively, cell B’ can also gain a communication capacity that was absent in cell B, for example the capacity to produce a synthetic ligand for communication. The model itself is general and any features of B’ cell can be different compared to B cells, e.g. proliferation, motility, division rate, etc. We note here that the cellular protein production and interactions are chosen to model contact-dependent signaling (also known as juxtacrine). Other signaling mechanisms, such as diffusion mediated patterning, can be implemented by choosing the appropriate differential equations.

Within this framework, cell rearrangements are non-linearly dependent on cell-cell signaling networks, adhesion preferences of the cells in the different states, and initial conditions. The amount of cell-cell contacts in fact changes over time as the systems restructure towards a stable configuration, which depends on pattern of cell adhesion preferences; but the stability of a given configuration also changes when the cell-cell adhesion properties of the cells change over time as a consequence of cell-cell signaling. This give rise to a non-linear system that cannot be treated completely analytically.

### Parametrization of the model

With the computational system at hand, we wanted to see if we could parametrize it such that it could describe available *in vitro* structures and networks from^19^. As detailed below, we first made decisions on basic parameters such as cell size and adhesion range with *in vitro* data, biological considerations, and feasibility constraints in execution time, then the rest of the more specific parameters were tuned by parameter scans and comparison with *in vitro* counterparts. In this way we identified a coherent ensemble of parameters that faithfully reproduce *in vitro* results. We note here that several different sets of numerical values for the parameters could give rise to the same cellular structures in CompuCell3D. For example, we chose 52 as cell-medium adhesion, which restricts the value of adhesion for more adherent cells in the 0-51 range; if we picked a different value for cell-medium adhesion, the precise numerical value for adhesion of the different adhesion molecule would be different.

Further details on the algorithm and parameters can be found in Methods, “*In Silico* L929 Cell Line Properties” and “CompuCell3D and the cellular Potts Formalism” and full lists of simulation parameters can be found in Table S1 for signaling and simulation and Table S2 for adhesion.

#### Baseline adhesion, cell size, deformability, and motility

Cells in the *in vitro* reference experiment are L929 murine fibroblasts; to create an *in silico* version, ISL929, we needed identify a parameter set that would produce biologically plausible cellular structures. The parameters that needed to be identified are highlighted in yellow in the CompuCell3D equations:

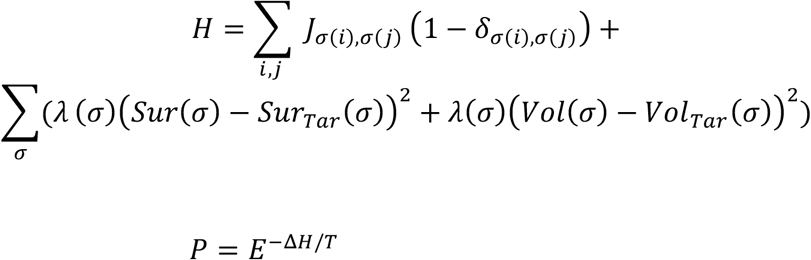

For the parameters for cell size cell shape, we first chose the target surface (Sur_Tar_) and volume (Vol_Tar_) of the cell to correspond to that of a sphere, as the cells are spherical before aggregating *in vitro*. If a cell is too small, then its movement is very volatile due to the small number of pixels that constitute it. On the other hand, a very large cell is slow and requires many MonteCarlo steps (mcs) to move, making it computationally demanding. With this motivation in mind, we set the cells to with a preferred radius of 3 pixels, corresponding to target volume and surface (VolTar and SurTar) of 113 pixels. Furthermore, we set the basal energy cost of deforming away from the target volume and area to be λ=2.2.

With these fixed parameter choices, we moved to identify parameters for temperature (T), which can be interpreted as a motility parameter^25,46^, and basal cell-cell and cell-medium adhesion via parameter screening. These parameters are crucial to define a multicellular system’s basal “mixability”, i.e. capacity of cells to rearrange in a spheroid. To parametrize T and baseline cell-cell and cell-medium adhesion, we first picked an arbitrary value for cell-medium adhesion, 52. The parameter choice for cell-medium adhesion sets the scale for the remaining model parameters, as cells that are not in contact with other cells are always in contact with medium, and they, for example, favor cell-cell only if the cell-cell adhesion is lower than the cell-medium. In this system, adhesion between cells contributes positively to the effective energy and, since the stochastic dynamics favor pixel re-arrangements that lower the effective energy, higher adhesion magnitudes correspond to a lower adhesion strength.

Then, we compared the behavior of an aggregate of 100 L929 cells *in vitro* with a similar aggregate of ISL929 cells *in silico* where parameter values of T and cell-cell adhesion were varied within a specified range (Fig. 3A-B). As visible from Fig. 3A, wild-type L929 cells weakly aggregate to a pseudo-spherical structure with “rough” edges, biologically indicative of weak cell-cell adhesion. *In silico*, a range of motility between 10 and 1000 and of cell-cell adhesion between 20 and 70 were tested. As shown in Fig. 3B, within 10,000 simulation steps (mcs), T=10 does not allow cells to move for any of the adhesion parameters. When T=1000, spheroid formation occurred only for cell-cell adhesion values of 20, whereas higher values for cell-cell adhesion (corresponding to lower adhesion strength) resulted in cells disintegrating from the spheroid, which is not observed *in vitro*. Moreover, with T=1000, cell shape was extremely distorted. At T=100, we obtained a spheroid with rough edges for cell-cell adhesion stronger than cell-medium (49 for cell-cell adhesion vs 52 of cell-medium adhesion – reminder that these values are inversely proportional to the strength of adhesion), similar to the *in vitro* phenotypes. These observations prompted us to use values of T=100 for basic temperature, 52 for cell-medium adhesion and 49 for cell-cell adhesion of cells. These values were used throughout the rest of the simulations presented in this paper.

When doing the adhesion parameter scan, we noticed that lower cell-cell adhesion values (higher adhesion strengths) resulted in larger aggregates, which was not consistent with previous observations *in vitro* that higher cell-cell adhesion results in more compact aggregates. To further investigate this, we cultivated aggregates of 100 L929 parental cells in U-bottom wells and compared their dimension to aggregates made of 100 L929 cells overexpressing E-cadherin protein, which leads to increased cell-cell adhesion strengths. We observed that the E-cadherin aggregates were indeed smaller than the parental ones *in vitro* (Fig. 3C). To recapitulate this behavior *in silico*, we did a parameter scan of the deformability parameter lambda over two adhesion values. As visible from Fig. 3D, a λ=5 does not allow cells to change shape at all, while λ=1 allows more highly adhesive cells (below 40) to form a compacted spheroid. When set at a value of 2.2, cells with adhesion above 40 formed larger aggregates, as seen *in vitro*. Based on this analysis, we chose to keep λ at 2.2 for parental cells, but to lower it to 1 for cells with adhesion below 40, such that they would mimic *in vitro* behavior and form smaller aggregates.

#### synNotch signaling for “2-layer” networks

To identify numerical values for the signaling equation (3.1), we use experimental data from a simple *in vitro* experiment presented in^19^. A population of receiver cells engineered with the synNotch receptor that activates a GFP target protein were set in contact with a population of sender cells expressing a cognate ligand. Then, fluorescent protein production is tracked over time (Fig 3E). The time-dependence of the normalized fluorescence provides the time scale of contact-dependent signaling. To mimic the signal induction dynamics in receiver cells *in silico*, a simulation was set up to track normalized protein production over time. Equal amounts of sender A and receiver B cells were seeded in two box-shaped sheets, with uniform contact between the two types. Levels of reporter gene in receivers were tracked over time for a total of 24,000 mcs. The signaling parameters β,κ,ε were heuristically adjusted (not shown) such that the simulation matched the experimental normalized fluorescence (shown in Fig. 3E) (numerical values for β, k, ε can be found in Table S1). These parameters capture dynamics specific to synNotch signaling in L929 cells and we used them as our baseline for all subsequent signaling interactions. Given the temporal dimension of the *in vitro* results, this gave us also a temporal translation between simulation time and real time of 1000mcs = 1h.

#### Adhesion in 2-layer networks

We then moved to identify numerical values for the adhesion weights (matrix J) of different adhesion molecules. In the *in vitro* experiments, different levels of E-cadherin (E-cad), N-cadherin (N-cad) and P-cadherin (P-cad) are used to modulate cell-cell adhesion. We first developed an initial relative hierarchy of adhesion strength based on available studies in the literature that pointed to differences in adhesion strength produced between cells expressing different adhesion molecules, or even different amounts of the same adhesion molecule^24^. In particular, E-cad, N-cad and P-cad are preferentially homotypical, but E-cad is stronger, and more promiscuous towards N- and P-cad whereas P- and N-cad are weaker and more selectively homotypical^47^. From this, we generated an adhesion hierarchy where, at similar levels of expression, E-cad produces greater adhesion strength between cells than either N- or P-cad; heterotypic E-cad:P-cad and E-cad:N-cad pairings are more favored than the N-cad:P-cad; and for any given cadherin molecule, low levels of expression produce less adhesion strength than higher levels of expression. With this general hierarchy of the relative strengths of different adhesion pairs we performed the following parameter screens to identify specific numerical values for adhesion strength parameters.

We started with identifying parameters for E-cad. To do so, we used experimental data from a training set structure that we call “2-layer network”: here, sender A cells activate receiver B cells to generate B’ cells that produce E-cad adhesion molecule. We knew from the *in vitro* data that if we started with 100 A cells and 100 B cells randomly mixed, in 24 hours we would obtain a 2-layered spheroid with activated B’ cells in the innermost layer, and A cells on the outside (Fig. 3F). *In silico*, we set up a simulation to mirror the *in vitro* experiment (Fig. 3G). Approximately 100 A cells expressing ligand_A were mixed with 100 B cells expressing receptor_A; when B cells acquire target protein units above the activation threshold (set at 5,263) they convert to B’ (Fig. 3H). We then ran 10 simulations for each value of B’-B’ adhesion between 25 and 50, and recorded screenshots of the resulting structure at 24,000mcs. The goal in the parameter screen is to identify parameters that achieve the maximal similarity to the *in vitro* picture, where activated, green B’ cells are found in the inner layer and A cells in the outer layer. As shown in Fig. 3H, for values of B’-B’ adhesion of 35 or higher, the sorting was either incomplete or the two types were randomly mixed. To get a quantitative measure of sorting, we also followed homogeneity index for both type A and B cells (Fig. 3I). We calculated a homogeneity index measure inspired from^10,48,49^, and defined in Eq S8 (Methods “Simulation quantifications” section) as the average fraction of surface area that cells of type X share with cells of the same type X over their total surface. The measure scores higher if the cells are uniformly contacting cells of the same kind, and lower if it has neighbors of a different kind. Given geometrical constraints, the highest value is around 0.8 for our cell numbers (as with a finite number of cells there will always be cells that do not have all their surface in contact with neighbors at all). Following this measure confirmed the impression from the visual inspection that: for values of B’-B’ adhesion higher than 35, the homogeneity of B/B’ cells was low, contrary to what is observed *in vitro* where B/B’ have highest homogeneity (Fig. 3I). These results constrained the value of adhesion of B’-B’ when they express E-cad to <35 and we picked the adhesion value for induced E-cad at 25 moving forward.

So far, we kept all the other adhesion to be at the parental level of 49, but we wanted to see if this was appropriate. To do so, we performed a similar screen for B-B adhesion values spanning between 25 and 50. In this case, if B-B adhesion is under 40, the sorting of the B cells precedes activation, and only B cells at the interface between B and A cells activate. This leaves the inner core of the B cells to the inactivated state, which is not what is observed in the *in vitro* counterpart. Therefore, B-B adhesion needs to be higher than 40, and we picked 47. We finally performed a screen for A-A adhesion values; this showed that low values of A-A adhesion results in formation of a blue core, and green poles at the periphery, which is not what is observed *in vitro*. Therefore, A-A adhesion values were constrained to >45 and we picked 49. With these parameters and starting with a mixture of approximately 100 A and 100 B cells, we consistently obtained 2-layer structures qualitatively similar to that of the *in vitro* results (Fig. 3J and see Fig. S1A for quantifications and more replicates, and MOVIE 1).

Collectively then, this screening allowed the identification of a set of adhesion values of B’-B’=25, B-B=47, A-A=49 (reminder that in CompuCell3D lower adhesion values correspond to higher adhesion strengths), that was able to recapitulate the fully sorted, 2-layer spheroid as in the *in vitro* experiment. The choice of the parameters is not univocal, any choice of the parameters in those ranges would have resulted in a 2-layer phenotype. We used these values for the rest of the papers for E-cad activation: induced E-cad – induced Ecad = 25; basal E-cad – basal-Ecad= 47, which is consistent with *in vitro* cells in a basal state having a leakiness of E-cad expression even in absence of activation^19^; this minimal difference between basal and parental (47 vs 49) is not sufficient to induce sorting (not shown here, but see fig. S2B.2 for an example where cells with 49 vs cells with 45 of adhesion do not sort).

Next, we used the parameter scan approach to parametrize cell type adhesion for N-cad and P-cad adhesion molecules. For induced N- and P-cadherin, we compared the parameter scan with the *in vitro* phenotype of a 2-layer network where B cells induce N-cad instead of E-cad. The phenotype *in vitro* is a less well-sorted 2-layer (Fig. S1B), which is similar to what is obtained *in silico* with values of B’-B’ of around 35. With this parameter, in fact, homogeneity of B/B’ is still higher than homogeneity of A cells, but lower than it was for E-cad expressing cells (compare Fig. S1A B/B’ green line with same line in S1B.4). Hence, for induced N-cad we chose 35 as adhesion value, and the same was then fixed for induced P-cad (given they are reported to have similar adhesion strengths^47^). Finally, another 2-layer structure from the training set allowed parametrization of heterotypic N- and P-cad adhesion, as well as constitutively-expressed P-cad. In this circuit, A cells constitutively express low levels of P-cad, whereas B cells are induced to express N-cad (fig. S1C). The resulting phenotype *in vitro* is that of a spheroid where A cells and B’ cells cluster together at different poles. Given the parameter scan in Fig. 3H-I, we thought we that, using A-A adhesion values in the range of 35-45 would allow us to achieve that; indeed, when we did simulations with A-A adhesion set to 43, we obtained *in silico* structures similar to the *in vitro* ones, where both A cells and B/B’ cells show similar homogeneity index values (Fig. S1C). This screening allowed us to identify parameters for adhesion values for E-, N- and P- cadherin.

#### Adhesion and signaling in 3-layer networks

More complex 3-layer structures can be generated *in vitro* with a more complex signaling network (Fig. 4A). In this network architecture, the B cells, when they receive the signal from A, transition to a B’ state that is able to communicate back to the A cells that can become activated to A’ cells. In this case, A, A’ and B’ cells have both the capacity to receive and to send a signal, a feature known as “transceiver” (Fig. 4C). This signaling logic is also called ‘back-and-forth’. This network was used *in vitro*^19^ to generate both central-symmetric 3-layered structures and also non-central symmetric structures, based on the choice of adhesion molecule. We chose to use the central symmetric three-layered structure as a training structure for parametrization *in silico*, and leave the non-central-symmetric structures for the test set.

The central symmetric 3-layer structure *in vitro* is generated as follows: A and B cell types have basal adhesion deriving from small amount of leakiness^19^, whereas A’ expresses low levels of Ecad (Ecad.Lo) and B’ expresses high levels of E-cad (Ecad.Hi). As shown in Fig. 4B, when approximately 200 A cells are mixed with approximately 40 B cells *in vitro*, A cells signal to B cells to induce Ecad.Hi, turn on reporter GFP, and form the center of the spheroid. Subsequently, surrounding blue A cells expressing blue fluorescence protein (BFP), are activated to A’ cells that turn on a red reporter (mCherry) and Ecad.Lo positive via signaling coming from activated B’ cells, resulting in a 3-layered structure with B’ cells in the center followed by A’ and then A cells at the outside.

To implement this *in silico*, A cells have ligand_A (filled circle) and can respond to ligand_B (green square) thanks to expression of receptor_B (squared). B cells have receptor_A (rounded) to respond to ligand_A, and in the B’ state can gain the capacity to send ligand_B (Fig. 4C). To parametrize the signaling for A and B cells, we proceeded as for signaling parametrization in the simper networks described earlier by comparing activation measures *in vitro* and *in silico*. For this more complex network architecture, the activation indices are defined based on cell type conversion from A to A’ and B to B’ as follows: *in vitro*, activation index is the normalized amount of GFP fluorescence (green) for signaling in cell B, and of mCherry fluorescence (red) for signaling in cell A (see methods “Simulation Quantifications”). *In silico*, activation index is the normalized ratio of activated cells over the total number of cells of the same type (See Methods, “Video analysis”). With these definitions we were able to compare *in vitro* and *in silico* signaling and identify parameters for signaling constants as described previously (numerical values for signaling parameters are in Table S1) (Fig. 4D). We notice that the signaling parameters are that allow for recapitulation of in vitro dataset are (slightly) different for cells A and B, even though they have a similar genetic network; we speculate that this could be due, *in vitro*, to different level of expression of the receptor or the transgene that makes signaling dynamics different in different cell types, hence parametrization of different cell types could require parametrization of signaling each time a new circuit is built. We used the identified signaling parameters for all subsequent simulations involving this kind of signaling.

For the adhesion values, the central symmetric 3-layer structure involves different expression levels of E-cad protein in different cells, e.g. low levels in A’ cells, and high levels in B’ cells. For the parametrization, based on previous work that established that changes in the expression levels of cadherins change adhesion strength^24,47^, we derived a hierarchy as in Fig. 4E. Based on this hierarchy, and starting from the previously identified value for Ecad-Ecad of 25, we chose to use Ecad.Lo-Ecad.Lo=40 and Ecad.Hi-Ecad.Hi = 20 (Fig. 4F).

With these parameters for signaling and adhesion, we then simulated the development of a system comprising around 200 A and 50 B initial cells. We observed that there was first induction of B to B’ cells, which then formed a green core. Then the B’ cells started to signal to the A cells to turn red (Fig. 4G, Fig. S2A and MOVIE 2). At the corresponding timepoints, we observed structures similar to that of the *in vitro* results (Fig. 4B), a 3-layer structure consisting of a green B’ cells core surrounded by concentric shells of red A’ cells, then blue A cells (Fig. 4G).

We wondered if, in the signaling scheme, reversion back to basal state played a role in this trajectory. To test this, we designed networks where there is no reversion, meaning that, once induced, cells cannot revert to basal state. Qualitatively and via homogeneity index, these two scenarios are not distinguishable in our setup (compare fig. S2A with S2C), suggesting that, *in silico*, the reversion might not play a big role for this specific trajectory.

Finally, we showed that signaling is necessary for 3-layer formation in the *in silico* model, similar to what was shown for the *in vitro* model. In the absence of signaling, there is no activation to B’ and A’, no formation of core(s), and no sorting occurring either qualitatively or quantitatively *in silico* (Fig. S2B).

This parametrization allowed us to identify values for all the target parameters that remain unchanged for the rest of the simulations. We then moved to test the quantitative capacity of the newly parametrized model to capture emergent properties of the system.

### Testing of predictive capacity of the model in the testing set

#### Parametrized model correctly captures emergent properties of in vitro developmental trajectories

With the model parametrized to capture the morphologies and activation dynamics, we wondered if the parametrized model would also reproduce other properties of the *in vitro* systems that were not used to calibrate the model parameters.

As a first example, we focused on the robustness of obtaining the target structures, focusing on the central symmetric 3-layer structure. Both the *in vitro* and the *in silico* systems have stochastic components. *In vitro*, the system forms a similar structure with one core 57% of the time experiments were performed (n=28 experiments^19^). In the *in silico* system, the stochasticity comes from the core algorithm in CompuCell3D, as the transition from one stage to the next is shaped by a probability distribution (Eq. 1). We quantified *in silico* the number of cores formed over repeated simulations of our parametrized model of the central symmetric 3-layer structure (n=30). The majority of the simulations yielded a 1-core structure (47%), some a 2-core (30%), and a minority a non-core structure (23%) (Fig. 5A). We compared this distribution to the distribution of morphologies reported for the biological system^19^, and found them similar (Pearson χ^2^=4.75, d.f.=2, P>0.09).

As a second example, we focused on changes in quantitative metric of morphology, and in particular the sphericity of the assembly. *In vitro*, we noticed that circularity of the structures evolves over time to reach a steady state by the end of the experiment, both for the overall structure and for the cadherin-expressing cells. To quantify these features, we used a standard circularity index in 2D that increases when the structure is more circular (see methods, “Quantification and statistical analyses”, Equation S9), and quantified it over time; results are in Figure 5B, blue and beige solid lines. To compare it with the *in silico* system, we defined a sphericity index *in silico* (eq. S7), and measured it over time. We plotted it together with the *in vitro* results and found that they generated qualitatively similar temporal evolution (Fig. 5B).

These data collectively show that the parametrized *in silico* system can recapitulate emergent properties of robustness and morphology evolution of the *in vitro* cellular system, properties that have not been used for the identification of the parameters.

#### Parametrized model predicts developmental trajectories with reshuffled adhesion

We now set out to test the predictive power of the parametrized based on changes to adhesion protein expression with reference to our available *in vitro* testing set. To do so, with the identified parameters, we ran simulations to see what the model predicts for the behaviors of multicellular systems with genomes that are obtained by changing adhesion values. Following the logic of one of the synthetic genomes in the *in vitro* test set, an *in silico* implementation had 2 cell types, A and B, and a back-and-forth signaling logic where A activates B to B’ and B’ activates A to A’. The adhesion features of the cells are basal for cells A and B, whereas A’ gains P-cad and B’ gains N-cad (Fig. 6A). We implemented this program *in silico* by giving A’ cells the value of adhesion that we identified previously for P-cad and similarly with B’ cells for the value of adhesion for N-cad. We then ran simulations with ~30 A cells and 30 B cells for 50,000mcs equivalent to 50hr of *in vitro* experiment. This gave rise to predicted structures that fell in two categories: one with one green B’ pole (90% of the runs), one with two red A’ poles (10% of the runs) (Fig. 6B, see also Fig. S3A for adhesion matrix, other sample structures and sorting index dynamics, and MOVIES 3 and 4 for runs with the 2 different phenotypes). When compared to the *in vitro* system, these structures display similar classes of morphology (Fig. 6C), with the one pole observed more frequently^19^.

We proceeded similarly for a second system (Fig. 6D), where the signaling logic is the same back-and-forth signaling between A and B cells, but now P-cad is constitutively expressed in A cells from the beginning, and signaling from B’ only changes color from blue to red. When we mixed approximately 160 A and 85 B cells for the equivalent of 34h, we obtained a structure with B’ green cells forming aggregates at polar positions in the spheroid, and the B’ aggregates are lined internally by A’ red cells; the inactive A cells stay in the center of the aggregate, since they express adhesion molecules from the beginning of the experiment (Fig. 6E, MOVIE 5 and Fig. S3B). The *in vitro* results obtained with engineered cells with similar genome resulted in a similar architecture (Fig. 6F).

These results show that our parametrized model can predict outside of the training set into new genomes obtained by changing the adhesion molecules produced by different cell types in the system.

#### Model predicts in vitro structures when initial number of cells are changed

We next explored the computational models’ capacity to predict final structures when initial conditions were changed. We ran simulations starting from different amount of A and B cells for three different systems.

For the first one, using the same underlying genetic architecture as in 6A-C, we increased the initial number of cells to around 90 for both A and B. *In silico*, increasing cell number resulted in bigger B’ green core, and formation of 2 red poles more systematically (Fig. 7B and S4A for homogeneity index and more replicates and MOVIE 6). In the *in vitro* experiment, it was noted that when more cells were used, the 2-pole phenotype occurred with higher frequency and the B’ green core was bigger (Fig. 7C) similar to what we obtained *in silico*.

For the second one, we returned to the genetic architecture underlying the central symmetric structure used to parametrize back-and-forth signaling (Fig. 7D). Instead of using the initial conditions from the training set (200A and 40B), we ran simulations with 160A and 90B. We obtained a thicker B’ central core and a thinner outer blue layer (Fig. 7E, Fig. S4B and MOVIE 7). When we looked at the *in vitro* data, it was noted that this initial cell combination led to a thicker B’ core as well (Fig. 7F).

For the third one, using the same underlying genetic architecture as in Fig. 6D-F, we decreased the initial number of cells to approximately 130A and 45B. *In silico*, the result of decreasing number cells resulted in formation of 2 green cores more systematically (Fig. 7H and S4C for homogeneity index and more replicates and MOVIE 9). In the corresponding test set *in vitro* experiment, a 2-pole phenotype is obtained (Fig. 7I) similar to what we obtained *in silico*.

These results show that our parametrized model can predict changes to the developmental trajectories when initial number of cells is changed.

#### The Model Recapitulates Synthetic Structures Generated by Lateral Inhibition Circuits Starting from Genetically Uniform Cell Populations

Having tested prediction capacity for changes in adhesion and changes in initial conditions, we wondered if the system could be predictive when the network itself was changed. To do so, we turned to another test set genome that involved a lateral-inhibition signaling network. Lateral inhibition networks are deployed during multicellular development, for example in the inner ear^28^, and entail a Notch receptor whose activation results in repression of its cognate ligand, Delta, in the same cell. This system has been modeled and studied extensively *in silico*, *in vivo* and *in vitro*. When deployed in architectures in 2D cellular lattices, lateral inhibition networks bifurcate to produce distinctive checkerboard patterning ^7,30,31,50–53^. In our computational system, thus far, we had parametrized signaling only for positive activations. Given that in the molecular logic, to obtain inhibitory signaling there is a swap of an inhibitory transcription factor in place of an activatory one, we hypothesized that our computational model could recapitulate inhibitory signaling by simply modifying our signaling equations (Equation 3) such that, S → - S (S is signal) and β→ - β (threshold of signaling). This would yield an inverse relationship between signaling and reporter production whereby low signaling receiver cells have high reporter production. We tested the inhibition version of the model on lateral inhibition by generating the following network in a monolayer sheet of cells: red A cell send and receive inhibition signals to/from neighboring A cells (Fig. S5A). In this setup, receptor activation decreases reporter level, so cell state A has target protein level above the activation threshold (we start from 7000), and can transition to A’ if its reporter inhibition is strong enough to make its reporter points fall below the threshold. In the A’ state the cells become green and loses capacity to signal. When we simulated development starting from red A cells in a single-layer sheet, we obtained the classic checkerboard pattern of lateral inhibition both when cells are regular (cells are not allowed to move, grow or divide) or irregular (cells are allowed to move but not grow or divide) (Fig. S5B). Importantly, since this was achieved keeping all the rest of the signaling parameters the same as for the other simulations described so far (adhesions are set at the parental level of 49 for all the cells), this shows that our parametrization can be extrapolated to simulate other known network contexts.

We moved then to see if this implementation of the lateral inhibition network could be used to predict the behaviors of spheroid morphogenesis based on lateral-inhibition and changes in cell adhesion. In the *in vitro* implementation, the logic of the signaling is that of lateral inhibition, but the adhesions are changed: A’ fate (green) increase E-cad expression, whereas A cells have basal adhesion capacity (Fig. 8C). When we computationally modeled this signaling logic (Fig. 8A) starting from approximately 100 A cells, initially some of the red cells become green, and then the green cells met each-other in the center of the aggregate. After 50h of simulated time, this results in a spheroid with both A and A’ cells, with the A’ green cells forming more preferentially the inside of the aggregate and never the outer layer, which was instead populated by inactivated red A cells (Fig. 8B and S8C for robustness and homogeneity index, and MOVIE 8). This phenotype is qualitatively consistent with the observation of the *in vitro* system, where a 2-layered structure is observed with green cells that adhere to other green cells, thereby forming a two-layer structure with a shell of red cells surrounding a green core^19^ (Fig. 8C).

Collectively these results show that our parametrized model’s predictive capacity can be extended to different network architectures.

### Model-based recommendation for genome for new structures

Finally, we asked if we could identify a network *in silico* that can give rise to a new structure not yet shown *in vitro*, a 4-layer central symmetric structure (Fig. 9A). We started from the observation that increasing the adhesive strength of the gray B cells in the two-layer model resulted in a homogeneous gray core with a thin layer of activated B’ cells on the edges (see Fig. 3H, bottom row, left most structure). This resembled a 3-layer structure but done with only a forward signaling between A and B cells. Thus, we hypothesized that if the B-B homotypic adhesion were increased to the inferred B Ecad.Hi adhesive strength, and if the inner B’ cells were allowed to activate the outer A cells, we might be able to create a 4-layered structure simply by reconfiguring the same synNotch circuits used in previous structures (Fig. 9B and Fig. S6A). This circuit configuration was tested with different initial conditions consisting of 2:1 mixed A and B cells at different number of cells, and allowed to run for 24,000mcs (equivalent to 24h *in vitro*) (Fig. 9C and Fig. S6B for more replicates). If the initial number of A cells is around 110 and B cells of around 70, there are not enough B cells to sustain the inner-most layer, resulting in loss of the inner-most layer, and forming a 3-layer-like structure at the end of the trajectory. If the initial number of A cells is around 1220 and B cells of around 620, however, there are too many cells impeding the formation of the homogenous inner core of (B/B’) cells, resulting in premature activation of (B/B’) cells. In this case, the final structure is composed of multiple, deformed inner cores, each surrounded by a homogenous layer of A cells. These results indicate that different initial number of cells, with the same circuit, could lead to different morphological outcomes. For the goal of a 4-layered structure with a single core, there exists a tradeoff between the initial number of cells and the resulting homogeneity of the layers in the desired structure. Our *in silico* experiments indicated that an optimal initial number of cells would be around 310 A cells and 170 B cells (Fig 9C and S6B). The recommended circuit would be as depicted in Fig. 9D, where A cells have ligand_A, and receptor_B that activates red and Lo.Ecad expression; whereas B cells have constitutive expression of Hi.Ecad, and receptor_A that activates ligand_B expression.

This shows that our computational system can generate recommendations for *in vitro* implementation of circuit for structures that have not been implemented yet.

## Discussion

A computational framework for the design of genetic networks for synthetic development would allow testing of the reachable morphogenetic space and allow identification of networks for user-defined trajectories and structures that optimize a certain parameter (e.g. burden on cellular machinery, number of cell-cell communication channels, number of cell types, etc.). This preimplementation optimization would provide access to a larger parameter space, allow identification of less intuitive solutions, and ultimately move the field towards the design phase. Here we provide a first step in that direction, by focusing on a paradigmatic example of developmental systems, where contact dependent signaling is paired with changes in cell adhesion in fibroblast spheroids.

Networks where cell-cell signaling changes mechanical properties of cells are a recurrent feature of development. The combination of signaling and morphological effectors has been shown to be at the core of complex developmental transitions: tissues are a complex system where cellcell signaling affects morphogenesis, and then morphogenesis feeds-back to influence signaling, to robustly generate complex multicellular structures^54^. In fact, the combination of signaling and morphological effectors has been incorporated in several computational models and shown to be able to replicate complex morphogenesis of embryonic transitions^1,9,23^. Given that the synthetic morphogenesis described in^19^ is one of the first synthetic systems that couples chemical and mechanical signaling for synthetic developmental trajectories in mammalian cells, it has attracted other computational description recently^42,55^, with different computational systems or objectives compared to ours. The system in^42^ describes a similar system in a cellular Potts model, with more focus on the underlying design principles and what can be learned about the logic of these kinds of networks. The system in^55^ is developed with predictive capacity and with some novel calibration structures, and relies on a new computational model for modeling cell movements that is distinct from cellular Potts. We think these efforts will collectively bring the field closer to the goal of rationally designing genetic networks for synthetic developmental trajectories.

Here we pursued one way to go about developing a computational system for design of genetic circuits of morphogenesis, using an available dataset to train and test our model for cellcell contact signaling and changes in cell adhesion. The computational system is designed to correspond to the synNotch receptor-based communication networks that was previously developed *in vitro*^19^. The modeling framework uses an available open-source platform, CompuCell3D, a cellular Potts based formalism that allows to model basic cell behaviors like proliferation, movement, and adhesion-based sorting^25^. Given that this computational system natively retains cell-cell neighboring relationship, which was originally used to compute the “entropy” of a multicellular system, we were able to use it to implement signaling that is dependent on how much contact there is between “sender” cells and “receiver” cell types. Signaling, in turn, is linked with activation in receiver cells to a different cell type that can be given different capacity for signaling and/or different levels of adhesion. This allowed us to create a modular backbone for implementing genetic networks based on contact-dependent signaling and changes in cell adhesion, which then generate synthetic developmental trajectories in cells.

We **parametrized** the model with *in vitro* data, using a subset of the complete *in vitro* dataset, what we call the training set. In particular, we parametrized signaling parameters for cellcell contact dependent synNotch signaling, and adhesion values for adhesion molecules of the cadherin family, using simple experimental setups. For the signaling, senders-receivers coculture experiments were used. For the adhesion parameters, simple, single-link networks were used, where signaling from A to B induces B to become B’, and where the different cell types can have different adhesion molecules. This setup allowed us to explore a parameter space and identify the *in silico* parameters that most closely mimicked the *in vitro* structures. The capacity of our system to match experimental results shows that the computational system can be tuned to represent synthetic developmental trajectories. Finally, the signaling for a more complex back-and-forth signaling network was parametrized with a simple example of that type of network *in vitro*. We used the simplest experiments to parametrize the model, mimicking a future pipeline where simple experiments would provide the baseline parametrization for predictive models that could identify potential circuit architectures for more complex implementations in cells.

Others have done parametrization in different ways: heuristic parametric tuning^42^ or machine-learning^55^. Our method was to design screenings of parameters in meaningful ranges, motivated by the biology and by computational considerations. It allowed us flexibility and exploration of a meaningful parameter space and was able to provide parameters with qualitative and quantitative matching. For more automated processes and extensions, the machine learning parameter estimation is an appealing future direction.

We then showed that the parametrized model can perform qualitatively accurate **predictions** of *in vitro* networks from a different subset of the *in vitro* dataset, what we call the test set. The test set had either adhesion molecules combination, initial number of cells, or network architecture that were different compared to the parametrization set. The capacity of our model to generalize outside the parametrization set suggest that our parametrization is a valid pipeline, at least for network designs with the same basic building blocks of signaling and effectors. Capacity to predict different initial condition is shown also by^42^. These efforts pave the way towards computational design of developmental trajectories.

One interesting aspect of our computational system is that it captures some of the robustness features of the *in vitro* system, for example in the formation of 1, 2 or multiple cores in the central three-layer structures. Noise and robustness are features of biological systems in general, and in particular for developmental systems^56,57^. Given our computational system is able to capture this important component, it seems it could be a useful resource to approach questions regarding whether certain network architectures are more conducive for buffering noise while still delivering the user-intended structures.

Finally, we showed how one can go about making a **recommendation** of a biological network for a user-identified phenotype. The fact that we have a parametrized model and know realistically implementable parameters, allowed us to do educated searches in a space that is smaller than the entire parameter space. These constrained searches could be helpful when trying to generate user-defined patterns or shapes starting with a limited toolkit of signaling and effector genes. Recent computational research showed that having access to a limited set of primitives is generative of a large number of shapes and patterns if you are allowing free parameter variations^58^. It will be interesting to see if this holds true in parametrized synthetic systems *in silico*, where you have access to the subset of parameters that are implementable. This would go in parallel with works where un-constrained approaches are taken^59^, which are helpful to identify needs for novel tools, like recombinases in the example. It will also be interesting to see how many of the structures obtained with the multi-recombinase approach can be deconstructed and re-implemented in our system with restriction to an implementable toolkit.

Our basic framework was built with the goal of designing artificial genetic circuits that would control synthetic developmental trajectories. Given this goal, we focused on “implementable” solutions, i.e. solutions that can be implemented with available genetic tools. To achieve this, our model modularly exploits the concept of cell types to signify cell transitions based on signaling. These transitions model acquisition of novel properties following cell-cell signaling events. In the work presented here, cell type transitions include changes in adhesion or signaling capacity. It would be rather straightforward in our model to extend to have the change in cell types signify changes in proliferation, motility, etc. This expansion would require initial parametrization experiments in simple setup, so that the user would know the parameters to use, for example, for the proliferation changes that can be executed with genetic controls.

Another interesting expansion for our system would be inclusion of other modalities of signaling such as signaling dependent on soluble ligands, bioelectrical signaling, as well as ECM mechanical and chemical signaling. Taking inspiration from what we achieved here with contact-dependent signaling, existing computational solvers for these other types of signaling could be incorporated to add or subtract points to cell type transition likelihood. This would be particularly interesting for soluble signaling, as morphogenetic signaling is another family of signaling that is predicated to underlie developmental transitions^54^ for which synNotch-based implementation has also been recently reported^60^. Modeling platforms exists to model changes of other morphogenetic currencies, like ECM-based, or bioelectrical in a tissue^61^, and one could imagine extension of our system to calculate cell type transition based not only on cell-cell contact signaling, but also on input from other computational engines.

Future directions for these computational efforts could be combination with artificial intelligence-based optimization either through evolutionary algorithms or machine learning. It has been recently shown that these could be used to generate morphologies that can then be recapitulated *in vitro*^62,63^. Algorithms could not only be trained to optimize parameters such as cell line, signaling network, and behavioral response, but could also incorporate subparameters such as: motility, proliferation, differentiability, juxtacrine and soluble morphogen signaling, mechanotransduction, adhesion, chemotaxis, and differentiation, to list a few. Numerous other recent advances in synthetic biology^33,64–71^ have made it possible to further control this process, facilitating synthetic reconstruction of complex native morphogenic processes towards enabling control over custom tissue development.

## Acknowledgements

The authors are grateful to the developers of CompuCell3D and the users of the help forum for their assistance in learning the program. The authors would also like to thank Marion Johnson and the other members of the Morsut lab, and members of USC stem cell department, Dr. Matt Thomson, members of the Thomson lab at Caltech for feedback that improved the work. The authors thanks Amanda Frataccia for her work on the scientific illustration of the project. This project was supported by a National Institute of Biomedical Imaging and Bioengineering R00 to LM (4R00EB021030-03), NSF RECODE from CBET-2034495, R35 from NIGMS R35 GM138256, along with a USC Department of Stem Cell Biology and Regenerative Medicine Startup Fund.

## Author Contributions

Conceptualization: C.L., L.M.; Methodology: C.L., L.M., S.S., G.C., J.C., C.C., D.Y.; Software: C.L., S.S.; Validation: C.L., S. S., G. C.; Formal Analysis: C.L., G.C.; Investigation: C.L., S.S., G.C.; Resources: L.M.; Data Curation: C.L., S.S., G.C.; Writing - Original Draft: C.L., L.M.; Writing – Review and Editing: C.L., L.M., S.S., G.C.; Visualization: C.L., L.M., S.S., G.C.; Supervision: L.M.; Project Administration: L.M.; Funding Acquisition: L.M.

## Methods

### KEY RESOURCES TABLE

| REAGENT or RESOURCE | SOURCE | IDENTIFIER |
| --- | --- | --- |
| Software and Algorithms |  |  |
| CompuCell3D (CC3D) v3.7.8 | <sup>25</sup> | RRID:SCR_00305 |
| Mathematica v11.3.0.0 | Wolfram Research | RRID:SCR_01444 |
| ImageJ v1.52a | <sup>72</sup> | RRID:SCR_003070 |
| JMP Pro v14.0.0 | SAS Institute | RRID:SCR_01424 |
| Excel v1808 | Microsoft | RRID:SCR_01613 |
| General Juxtacrine Signaling Model (GJSM) in | This paper | N/A |

### CONTACT FOR REAGENT AND RESOURCE SHARING

Further information and requests for resources or code should be directed to and will be fulfilled by the Lead Contact, Leonardo Morsut.

### COMPUTATIONAL METHOD DETAILS

#### CompuCell3D and the cellular Potts Formalism

We implemented our model in CompuCell3D (CC3D) v.3.7.8 ^25^, a modeling software that allows simulation of cells and their behaviors using the cellular Potts formalism. By itself, CC3D contains numerous built-in features for replicating *in vitro* cell behavior, several of which we utilized either directly or adjusted via CC3D Python v.2.7.13 scripting according to manual v3.7.9. In our model, we incorporated default features from CC3D such as surface area constraint, volume constraint, cell division, adhesion, cellcell surface contact, and cell types. We implemented custom cell motility, cell growth, and cell signaling, as described below and in subsequent sections.

We defined cells as multi-pixel entities in 3D that physically act by performing “pixel copy attempts” over simulation time steps (monte carlo steps, mcs). Performing “pixel copy attempts” effectively moves and changes both cell geometry and position over time. These pixel copy attempts succeed probabilistically, determined by the Boltzmann acceptance function, P=e^-ΔH/T^, where P is probability of attempt success, ΔH is change in total effective energy of the system from all attempted pixel copy attempts at the mcs t, and T is the cell motility ^25^.

Effective energy (H). Because we incorporated surface area constraint, volume constraint, and adhesion, our total effective energy H at a given mcs t therefore takes the form,

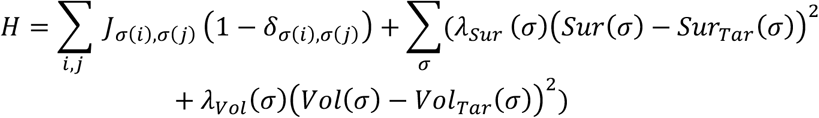

as described in ^3^. The terms σ(i) and σ(j) denote the identity of the cells occupying pixel sites i and j separately, with the Kronecker Delta limiting inclusion to only the cell interface. J is a matrix that contains the contact energy between cells of different identity. λ_Sur_ and λ_Vol_ constrain deviations of a cell from the ideal surface area Sur_Tar_ and Vol_Tar_, hereafter referred to as target surface area and target volume, respectively.

The numerical values in J control adhesion in cellular Potts. The values in J represent a stability index: lower J makes for a more stable state, which is then how you achieve stronger adhesion. Conversely, a higher J leads to weaker adhesion.

#### Generalized Juxtacrine Signaling Model (GJSM)

Juxtacrine signaling is the method employed to achieve the known synthetic structures. For a generic signaling ligand whose expression was constitutive, constant, and unaffected by signaling, we describe the total **ligand level**, L, on a cell’s surface by the equation

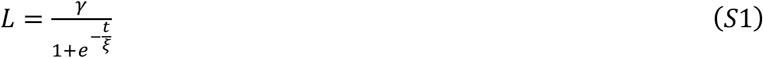

where t is the given time in mcs, while γ, and ξ are constants. We chose this equation because of its simplicity. It could be generalized to represent steady state ligand level on a cell’s surface, recovery of surface ligand level from trypsinization, and experimental conditions such as ligand induction via tetracycline from a drug-controlled promoter (e.g. Tet On).

Then, a receiver cell in contact with the sender cell would change its **target protein level**, R, by the differential equation

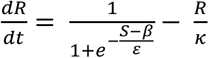

where β, ε, and κ are constants, whereas S is the signal strength. We chose this form for several reasons. First, parameters have intuitive interpretations: β controls sensitivity to S, ε modulates magnitude of S and β, and κ represents the standard linear protein decay rate constant commonly employed in biological models. Secondly, these parameters have kinetic/biological interpretations, due to the logistic function’s intrinsic relation to the Hill function^73^. Lastly, this form of the logistic function is easily tunable and well behaved, due to its monotonicity from negative infinity to positive infinity and bound between 0 and 1. This tunability is not as easily achievable with the Hill function, where odd or fractional Hill constants lead to the existence of singularities.

In the case where the target gene is a ligand itself, we use equation (2) to calculate ligand levels.

The time-dependent evolution of the reporter, apart from the parameters, depends on **signal strength** S; this reflects the biological fact that the promoter of the target gene is under the control of the receptor in juxacrine signaling. Signal strength is itself affected by: the number of receptors on the receiver cell (Ω), the number of ligands on contacting neighboring cells (L), the surface contact area between receiver cell and its neighbors (Φ). Because these factors evolve over time, S is therefore a morphological dependent and time dependent function that evolves according to structure’s spatial organization.

The following describes how we take into consideration the shared surface area to compute S. We consider a single receiver cell σ; first we need to identify which of its neighbors can signal with it. The different cells are assigned different types, according to whether they can signal (ligand expressing) or can receive (receptor expressing) or both. These cell types are indicated as A, A’, B and B’ in the text. For example, we have a cell σ of type A, that express receptor r_A_, that can be activated by ligand l_A_. This allows us to identify the neighbors of sigma, by looking through the list of all neighbors of sigma and identifying the ones that are of a type that bear ligand l_A_.

Then each different ligand/receptor interaction is treated identically regardless of the specific mechanism (e.g. if it models anti-GFP/GFP or anti-CD19/CD19).

Receiver cells has receptors on its membrane, quantified by Ω_σ_. The neighbors have ligands on their membrane, quantified by L_i_. To compute the amount of receptor-ligand interactions that can happen when receiver + cognate sender cells are in contact, we need to calculate the amount of receptor and ligand that are present on the surface of contact.

To do that we first define the portion of contact surface for sender (SNi) and receiver (sigma):

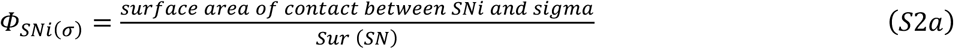

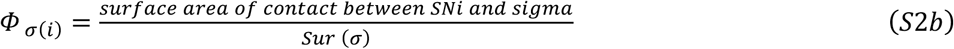

These are now multiplied for the total amount of ligand (or receptor), to obtain the amount of ligand (or receptor) that is available at the area of contact.

Available ligand = Φ_SNi(σ)_ * L_i_

Available receptor = Φ_σ(i)_ * Ω_σ_

With these two values we can calculate the value of S_σ_ as follows:

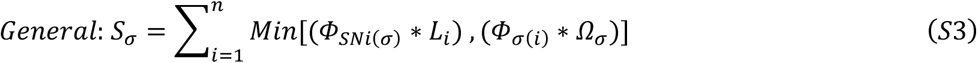

where n is the total number of cells that are currently in contact with sigma and that can engage in signaling with sigma, i.e. produce the ligand for which cell sigma produces the receptor.

This results from assumption of:

a 1-1 stochiometry of 1 ligand activating 1 receptor, given the biochemistry of the signaling;
homogeneity of ligand and receptor on the cell’s surface.

This is in general; in the majority of the simulations that we describe in the results, we employed a simplified version where the receptor is considered to be in excess, as we do not have evidence to think otherwise. In this case, the only part of (5) that is determining the amount of signal S is given by the ligand, so S takes the form of:

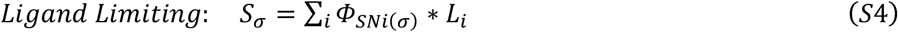

To implement signaling-inducible behavioral response, we use a state-transition model. We borrow notation from physics; cells of each genotype, if excitable, bear a ground state and an excited state or even multiple higher order excited states. The transition between states is regulated by the target gene activation, which is regulated by the cell-cell signaling. Cells in different states can have different properties such as color/state, adhesive properties, deformability properties, signaling/reception capacity (Fig. 2F). This quantized representation of cell behavior has been applied, though not with this notation, in other models^3,4,45^. In this way, the signaling can induce behavioral changes in the cells that receive the signal, generating a highly non-linear system of interacting agents.

Because the reference experiments primarily focus on signaling inducible adhesion with reporter, we utilize two states per genotype, ground and excited, in the biological replication simulations.

The excited state bears a different color from the ground state, reflecting signaling induced reporter expression. Adhesion matrix J can be defined for the different states, to mirror changes depending on adhesive strength and binding specificity that the cadherin types in the *in vitro* counterpart express upon sufficient signaling (see Table S1). It is also possible for a cell to fall from the excited state to the ground state due to loss of signaling; falling under the transition threshold will move an excited state cell to the ground state, reverting color and excited properties.

#### *In Silico* L929 Cell Line Properties

##### Cell division

*In silico* L929 (ISL929) cells consist of multiple pixels and starts with a target radius (TR) randomly chosen using a Gaussian distribution (μ=3.0 pixels, σ=0.5 pixels). This TR is then used to calculate the target surface area (4πr^2^) and target volume (4πr^3^/3) for each cell, as *in vitro* L929 cells adopt a spherical shape at the beginning of experiments ^19^. Each cell then undergoes growth by experiencing net positive increase in TR from small positively skewed uniformly distributed fluctuations in TR. Target surface area and target volume thus increase slowly over time. Upon reaching a threshold volume, 2*4πμ^3^/3, the cell then undergoes division, resulting in the original cell and a new cell. The original cell is subsequently reassigned a new TR from the above Gaussian distribution and both target surface area and target volume are recalculated. The new cell is assigned the same post-division parameters as the original cell, modelling completely symmetric cell division. All the parameters and state variables are inherited by the two daughter cells from the parent cell. These choices result in roughly doubling time of 24,000 mcs (equivalent to 24 hours, the estimated doubling time of L929 cells ^19^). For example, this resulted in a ratio of cell number at t=24h vs t=0h of

2.07+- 0.10 (n=10) for simulations in Fig. S1A;
2.02 +- 0.08 (n=10) for simulations in Fig. S1B.5;

And ratio between 20h and 0h of:

1.73+-0.08 (n=30) for Fig. 4B

We note that, due to stochastic nature of growth, cell death is also possible within this model.

##### Cell adhesion

*In vitro* L929 mouse fibroblasts weakly adhere to one another under ultra-low attachment suspension conditions ^19^ thus we designate our basal, parental ISL929 cells to have a relatively high J to one another and a slightly higher J to the medium, resulting in the formation of weak aggregates in medium. As a result, these ISL929 cells also bear high motility, again similar to *in vitro* L929 ^19,74^.

##### Cell deformation

Cells that express adhesion proteins and adhere to each-other in vitro, deform markedly, and lose their rounded morphology ^19^. In CC3D, a way to change deformability of cells is through modulating parameters λ and in Eq (1) for H, with lower values corresponding to higher deformability. Cells with an adhesion matrix value of at least 39 (i.e. 0-39 range) (see Table S1), λ and λ were set to 1.0. Other cells had λ and λ set to 2.2.

##### Cell motility

Cell motility is defined in CC3D via the parameter T. Biologically it is known that cell adhesion to environment is complexly linked to cell motility, and adhesion effects on motility vary widely between different adhesion proteins and cell types ^75–77^. In general, although clearly not all-encompassing, the adhesion abstraction is that strong cell adhesion to environment tends to decrease cell motility^75,77,78^.

For our purposes, in the *in vitro* L929 system, we noticed that the cell motility is rather similar across different cell types and different adhesion, with some minor differences between adherent cells (slightly lower motility) compared with non-adherent cells (slightly higher motility). We therefore defined motility as a sum of a constant T0 plus function of a cell’s environment (neighboring cells and medium), so that higher adhesion results in lower motility; in this way, different cells can have different motility. Each cell’s individual motility T_σ_ is:

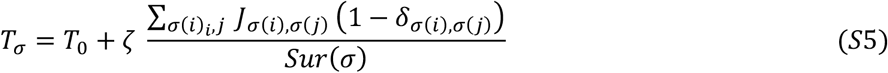

This formula iterates over each neighboring cell pixel and medium uniquely, and ultimately T is determined only by the type of the focal cell, the types of the neighbors, and total contact with medium. Categorizing environment by cell types and medium instead, accomplished in CC3D via cell-cell surface contact feature and cell type index, we obtained a computationally simpler approximate formula, which is the one that we use in our simulations:

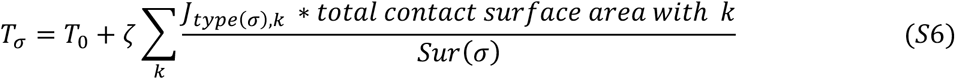

T_0_ is a constant representing basal cell motility, ζ a constant representing how effectual adhesion is at attenuating motility, and k denotes “cell type” (can be either a cell type or medium). This T allows each cell to sense its adhesivity to local environment, decreasing motility if adherent to neighbors and restoring motility when exposed to non-adhesive conditions.

T_0_ and is ζ are kept constant throughout after the initial parametrization in main Fig. 3B.

#### Simulation Lattice

Simulation lattice. At the center of a 100×100×100 lattice, we seeded a mixture of (A) and (B) cells as a radially symmetric blob to maintain a consistent initial cell aggregate shape while also maintaining a similar cell total and ratio to that of the reference experiment. For the setup in “2D” (Fig. S5B) we used a 100×100×5 pixel cell monolayer (~400 cells).

### QUANTIFICATION AND STATISTICAL ANALYSES

#### Simulation Quantifications

##### Sphericity index

(B’) green cells were visualized in 3D to determine core amounts and counted for each simulation at the endpoint. Sphericity was measured over time, both for excited states and over all states (Fig. 3d), using the formula ^79,80^

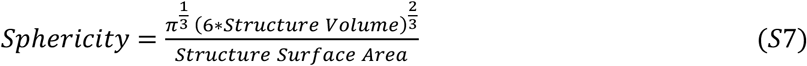

We roughly rescaled the sphericity by dividing by 0.48 to compensate for the cubic nature of the voxels. We measured activation timescale by measuring the number of (B’) and (A’) cells present per timestep and normalized each to 1 maximum.

##### Homogeneity index

We were interested in the spatial patterning of different cell types in these multicellular structures over time, thus we developed and quantified homogeneity index Ψ per cell type X, calculated according to the formula below

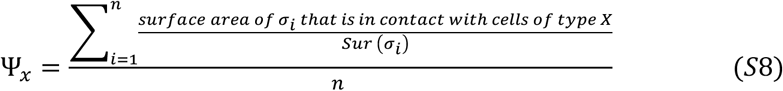

Where the sum is taken only for cells of type X that are in contact with other cells of type X, and n is the total number of cells of type X in contact with cells of type X.

This measure ranges from 0 to 1, with 1 indicating maximal homogeneity, and is similar to sorting measurements employed in other studies ^10,48,49^. By focusing only on cells that have neighbors of the same kind, this measure focus on clusters and not on isolated cells.

This measure hence depends also on the number of cells present, given that if you only have 2 cells for example, even though they are very homogeneous, less than 100% of their surface will be occupied by the neighbor.

This measure can be generalized for more than one cell type, by considering for example two cell types together. For example, in our simulations, when we have a genotype B that gives rise to two cell types (B basal and B’ excited), we may be interested in measuring homogeneity index for B and B’ combined. To do so we calculated Ψ_B,B’_ by defining type X as {type B or type B’} in the above formula. If desired, this measure can be simply extended to the ground and excited states of each genotype as well, Ψ_A_, Ψ_B_, Ψ_A’_, Ψ_B’_, or condensed as desired, Ψ_A,A’,B,B’_, making it possible to distinguish the effects of different behaviors on morphogenesis. This measure can be applied to many different morphologies, beyond fixed lattices ^49^ and spherical morphologies.

##### Core Distribution

We counted the number of cores per structure by visualizing only (B’) green cells in 3D at the endpoint of each simulation. To determine whether a core was a single core or double core, we visualized the endpoint simulation (B’) green cells from multiple perspectives to prevent viewpoint bias. This allowed us to determine whether there was truly a single core or multiple cores of similar sizes, with the latter generating 2-cores or 3+ cores (counted as “other”in Fig. 5A). Small cell stripe connections between cores were negligible and therefore counted based on the cores. Structures that did not appear core-like (i.e. large cell stripes) were counted in the “other” category. See Fig. S7 for examples of core counting.

##### Activation index

the activation index *in silico* is defined as the normalized ratio of activated cells over the total number of cells of the same type, i.e. #(A’) / Maxt[(#(A’)] and similarly for (B) and (B’) cells.

#### Video Analysis

*In vitro* data was either provided in the reference paper or obtained by analyzing the supplementary video for the counterpart structure from the reference experiments ^19^.

##### Circularity index

The video was split into constituent frames using Mathematica v11.3.0.0, then circularity analyzed by drawing a region of interest around the structure using ImageJ v1.52a, both in bright field (all cells) and merged color field (activated cells only), and data collated in Microsoft Excel v1808. Circularity was then calculated using the classic equation

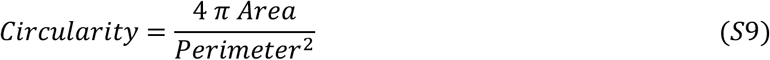

This measure ranges between 0 and 1 and the closer to 1 is indicates it is closer to a perfect circle.

##### Activation index

To estimate how fast cells activated over time, we color separated the green, red, and black merged image portion of each frame by green and red to generate two sets of frames, one for green and one for red, representing respectively the activated cells of (B) and (A). We then converted these frames into binary images using the MorphologicalBinarize function in Mathematica, replacing pixels with an intensity above 0.1 with pixels of intensity 1. This threshold value was minimally low to remove non-cellular background fluorescence and prevent biasing activated cell detection. Binarization additionally facilitated comparison by splitting *in vitro* cells into discrete states. Totaling the pixel intensity for each frame of each set estimates activation per timepoint for (B) and (A). Cellular background fluorescence, due to a few cells beginning with some green/red^19^ was removed by subtracting the minimum background fluorescence of the time series. Using the minimum helped negate cellular background fluorescence with again minimal biasing of activated cell detection.

#### Statistical Analyses

Sample sizes are given in the text and/or figure caption. Statistical tests were performed in JMP PRO v14.0.0 with a significance level of 0.05. We performed a chi-squared analysis for our core distribution analyses (Fig. 5A and Fig. S2C.4). Appropriate test was chosen according to data type and assumptions tested by residuals analysis. We report and show mean ±s.d. for all measures.

### DATA AND SOFTWARE AVAILABILITY

All simulations were performed in CompuCell3D v3.7.8 with custom scripts coded in Python v2.7.13. General source code and specific codes for the structures presented in this study are available at [https://github.com/lmorsut/Lam_Morsut_GJSM]. Dataset for the structures are available upon request.

#### Supporting Information

Supplemental Figures 1-7; Supplemental Figures Legends; Movies 1-9; Movies Descriptions; Table S1 with signaling and simulations parameters; Table S2 for adhesion parameters.

