## Supplemental Figures for "A parametrized computational framework for description and design of genetic circuits of morphogenesis based on contact-dependent signaling and changes in cell-cell adhesion"

#### Fig. S1, related to Fig. 3

**(A)**

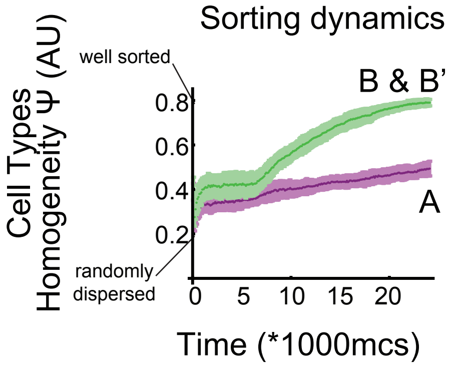


Adhesion Matrix

| | A | B | B' |
|---|---|---|---|
| A | 49 | 49 | 49 |
| B | | 47 | 42 |
| B' | | | 25 |

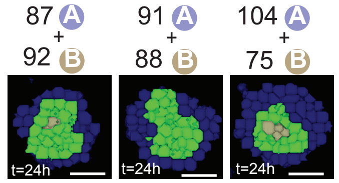


**(B)**

**(B.1)**

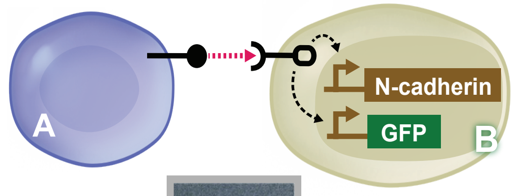


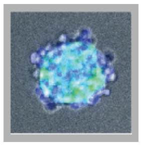

Reproduced from Toda et al., 2018

**(B.2)**

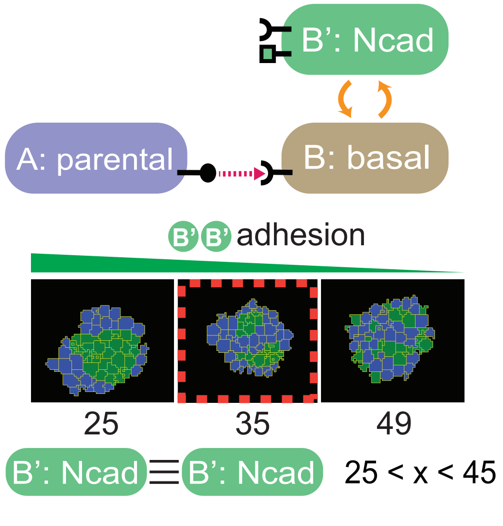


**(B.3)**

Adhesion Matrix

| | A | B | B' |
|---|---|---|---|
| A | 49 | 49 | 49 |
| B | | 47 | 42 |
| B' | | | 35 |

**(B.4)**

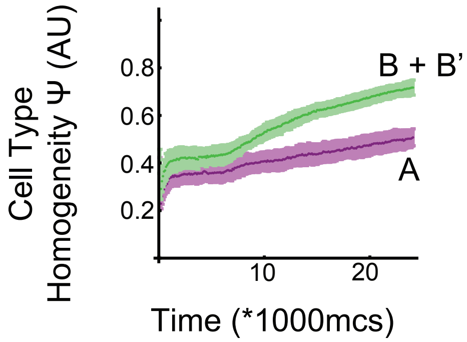


**(B.5)**

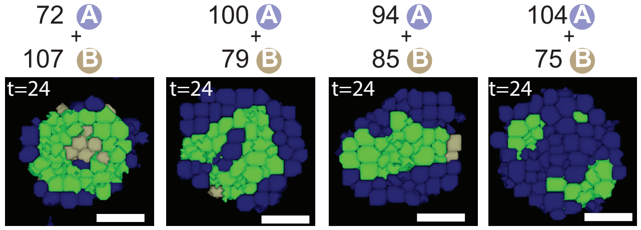


**(C)**

**(C.1)**

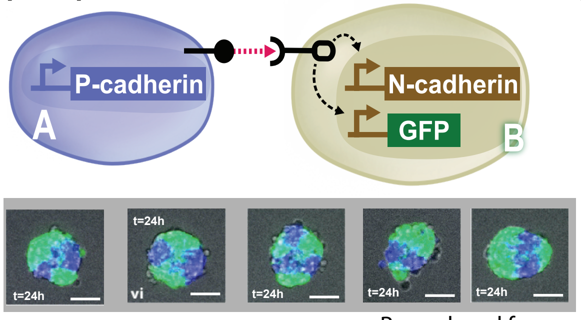


Reproduced from Toda et al., 2018

**(C.2)**

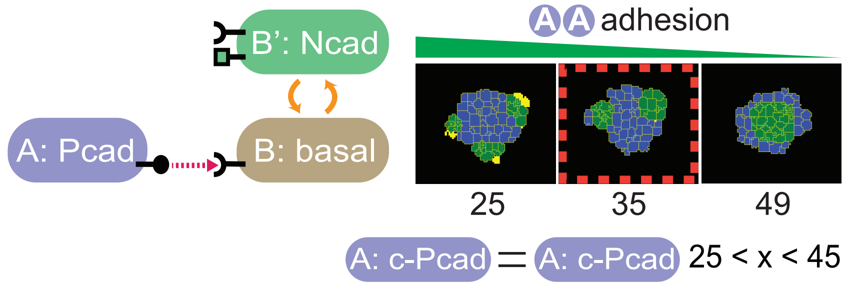


**(C.3)**

Adhesion Matrix

| | A | B | B' |
|---|---|---|---|
| A | 43 | 49 | 49 |
| B | | 47 | 42 |
| B' | | | 35 |

**(C.4)**

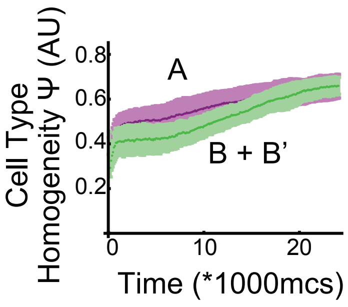


**(C.5)**

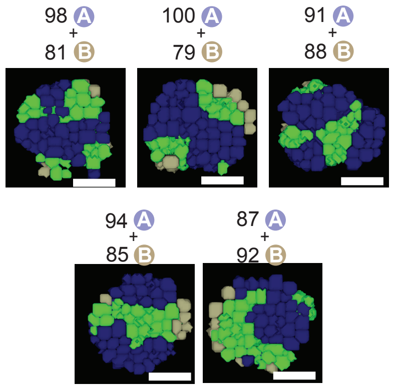

**Fig. S2, related to Fig. 4**

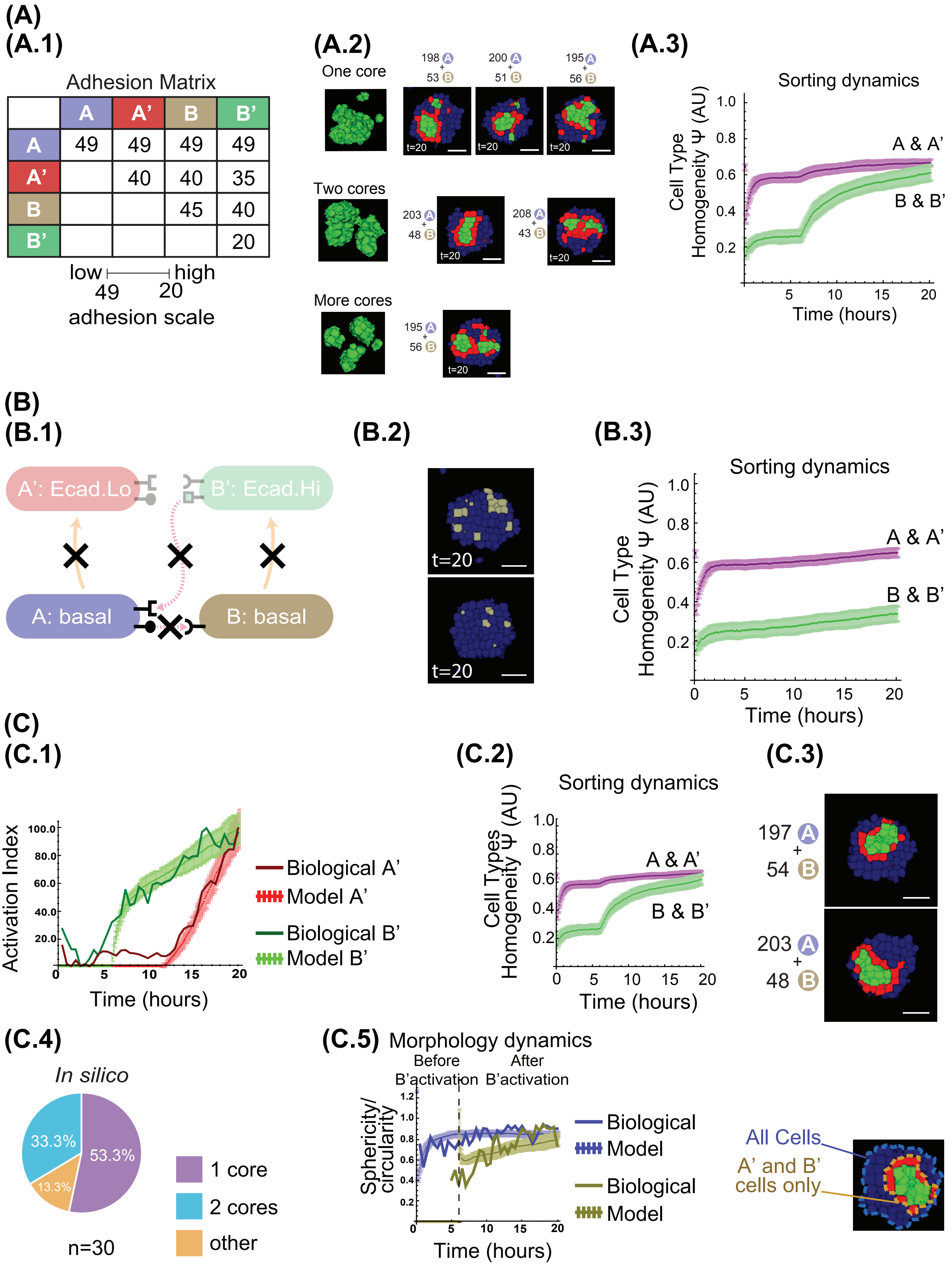

#### Fig. S3, related to Fig. 6

**(A)**

**(A.1)**

Adhesion Matrix

| | A | A’ | B | B’ |
|---|---|---|---|---|
| A | 47 | 42 | 49 | 49 |
| A’ | | 35 | 49 | 49 |
| B | | | 47 | 42 |
| B’ | | | | 35 |

**(A.2)**

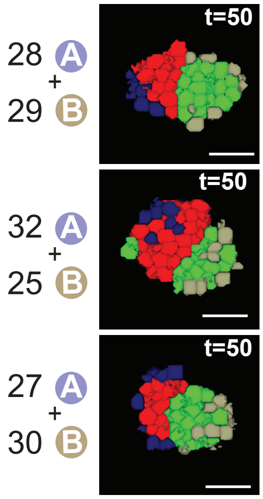


**(A.3)**

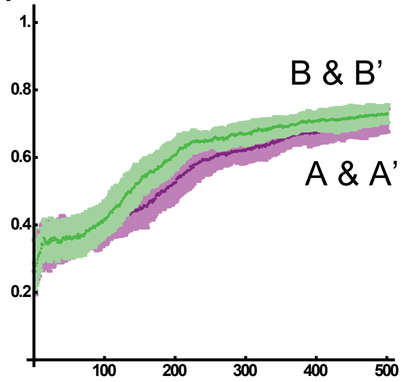


**(B)**

**(B.1)**

Adhesion Matrix

| | A | A’ | B | B’ |
|---|---|---|---|---|
| A | 43 | 43 | 49 | 49 |
| A’ | | 43 | 49 | 49 |
| B | | | 47 | 42 |
| B’ | | | | 35 |

**(B.2)**

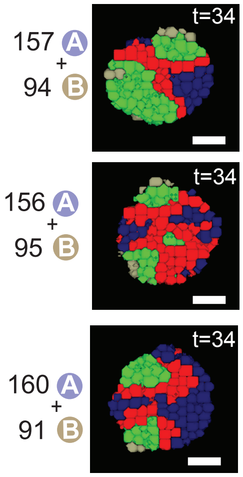


**(B.3)**

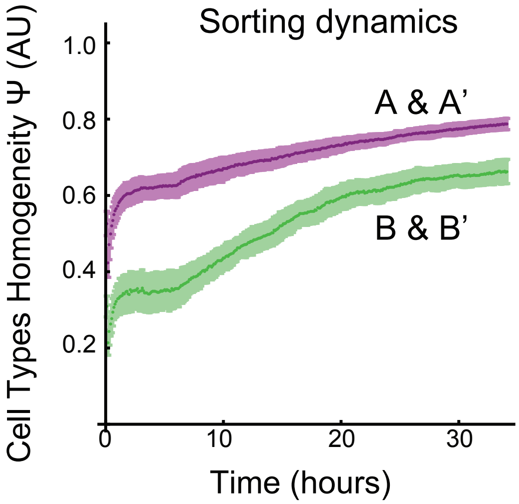


**(B.4)**

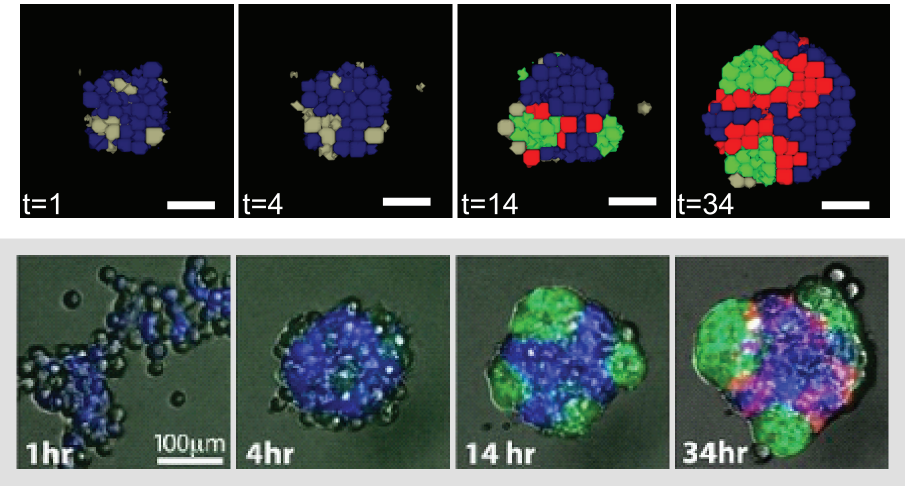


Reproduced from Toda et al., 2018

### Fig. S4, related to Fig. 7

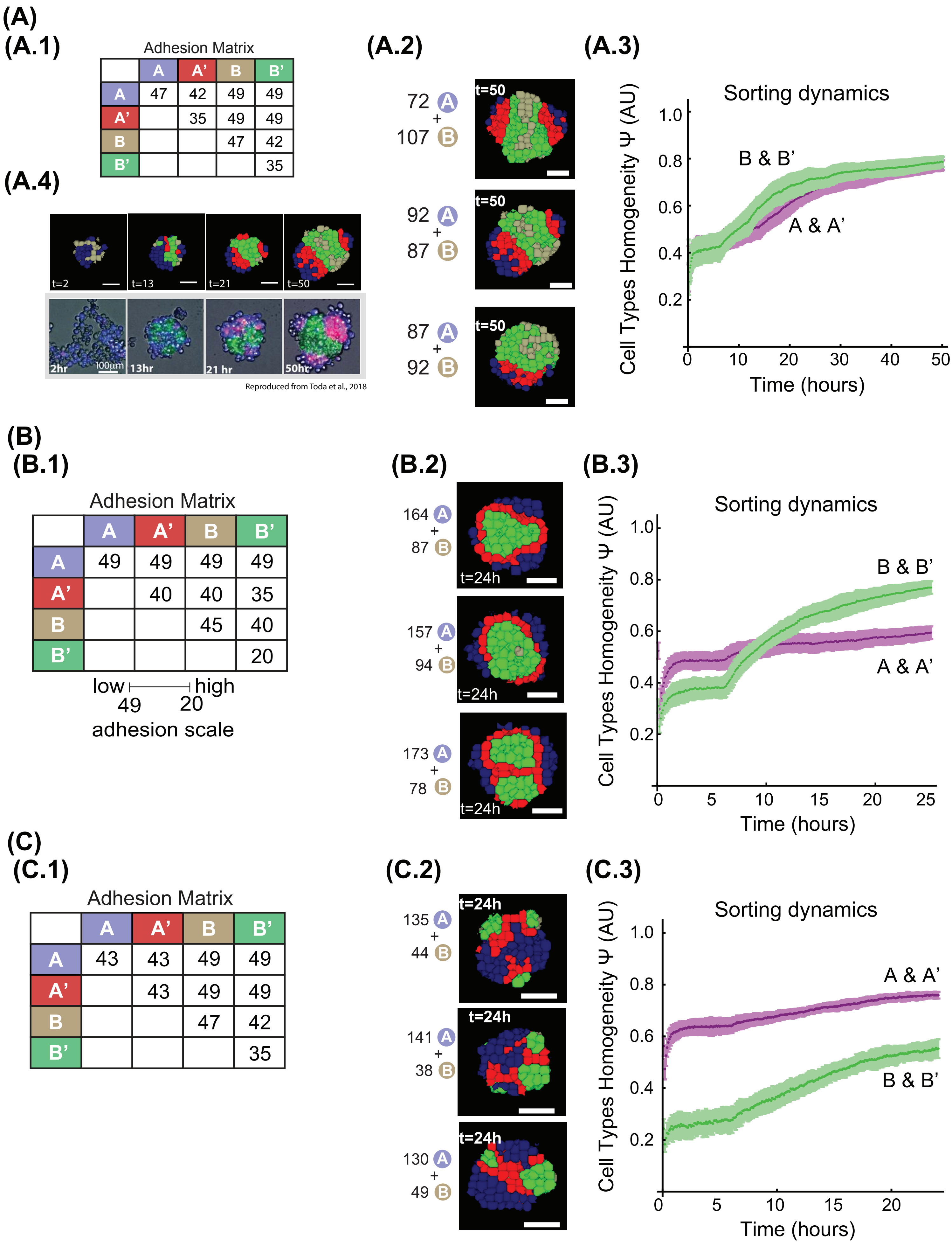

#### Fig. S5, related to Fig. 8

**(A)**

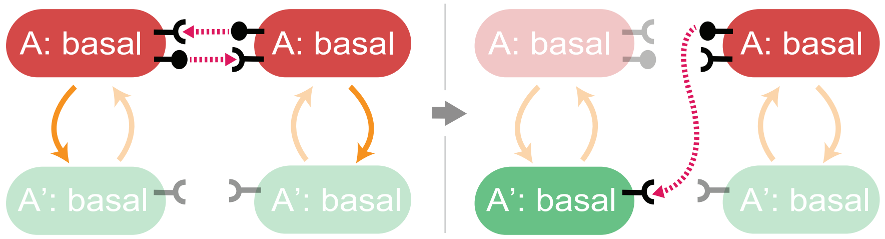


**(B)**

cell shape regularly constrained | cell shape irregular

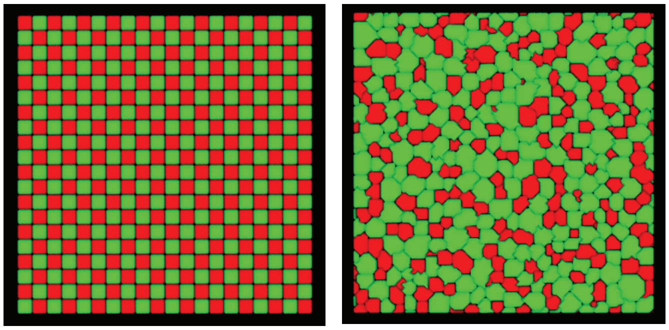

**(C)**

**(C.1)**

Adhesion Matrix

| | A | A' |
|---|---|---|
| A | 47 | 42 |
| A' | | 25 |

**(C.2)**

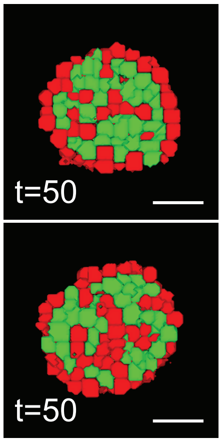


**(C.3)**

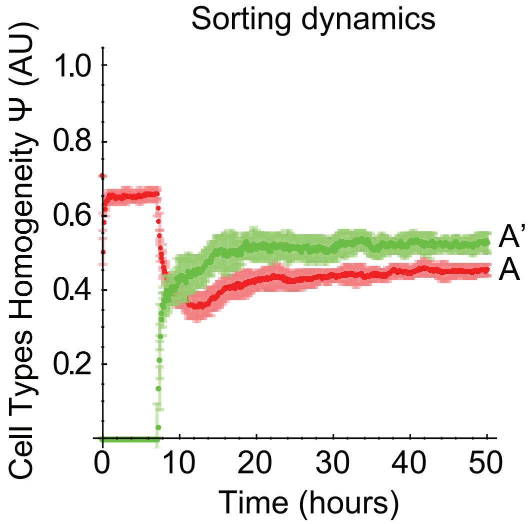

#### Fig. S6, related to Fig. 9

### (A)

Adhesion Matrix

| | A | A’ | B | B’ |
|---|---|---|---|---|
| A | 49 | 49 | 49 | 49 |
| A’ | | 40 | 35 | 35 |
| B | | | 20 | 20 |
| B’ | | | | 20 |

low ⊢—⊣ high
49 20
adhesion scale

### (B)

109 A
+
68 B

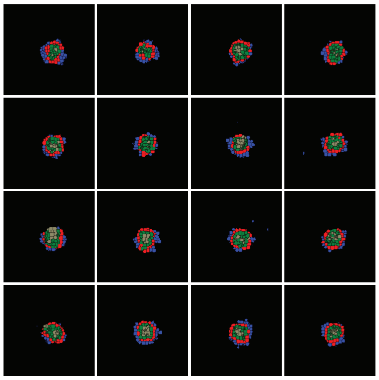

312 A
+
171 B

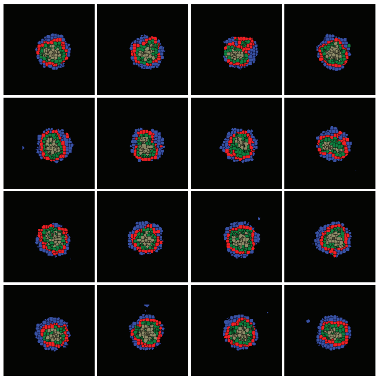

1219 A
+
618 B

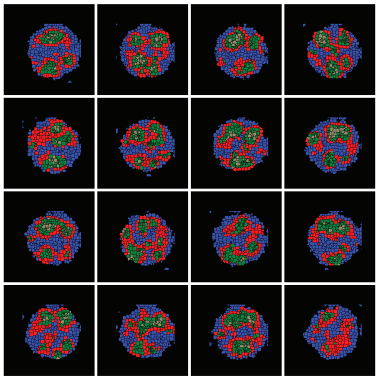

### Fig. S7, related to Methods

**Rules:**

- Score visualing only the 3D green cells to prevent perspective issues (we don't need to choose a plane to judge from, plus we can rotate the structure so nothing is hidden from our view)

- Score 1 per core, if multiple cores exist, count the cores that are similar in size. Bascially exclude the small irregularities.

- Some cores will bear a weak/striped connection between other core like structures. We will estimate conservatively using the classification below.

  twin connected core (TCC) = 2 cores
  triple connected cores (TCP) = 3 cores
  WKY = cannot confidently classify, count as minor like in Toda notation

**1 core examples**

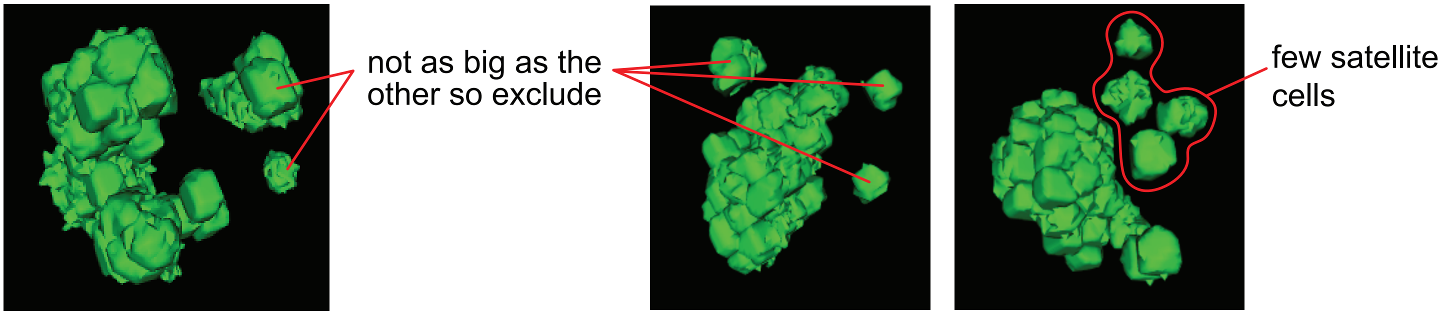


**2 core examples**

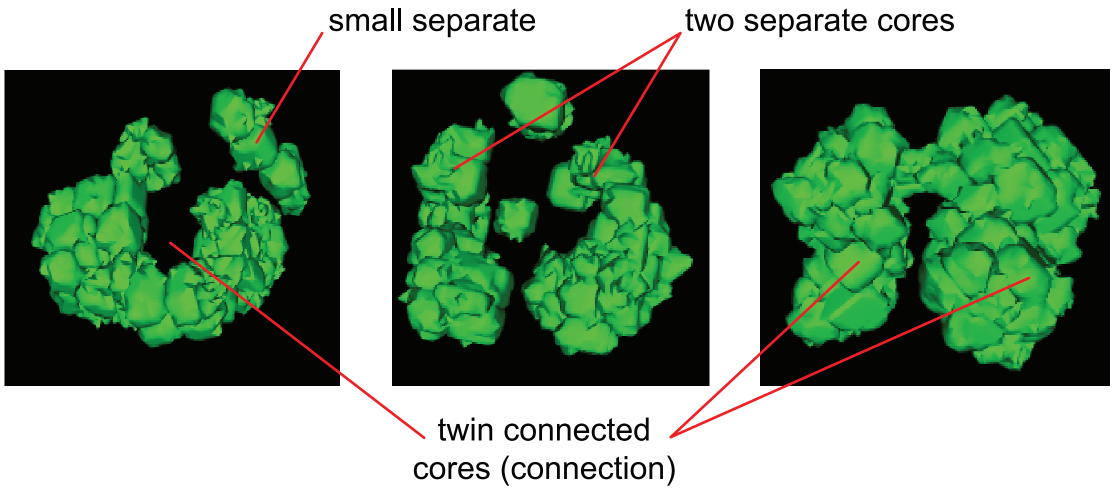


**3 core examples**

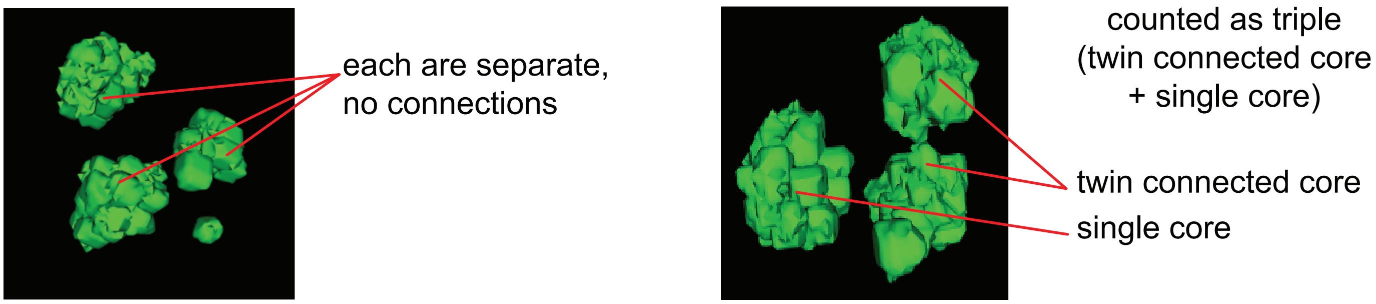
