## Supplementary material for "A parametrized computational framework for description and design of genetic circuits of morphogenesis based on contact-dependent signaling and changes in cell-cell adhesion": Table S2

| | Induced **Hi.Ecad** | Basal **Hi.Ecad** | Induced **Ecad** | Basal **Ecad** | Induced **Lo.Ecad** | Basal **Lo.Ecad** |
|---|---|---|---|---|---|---|
| Induced **Hi.Ecad** | 20 | 40 | | | 35 | 49 |
| Basal **Hi.Ecad** | | 45 | | | 40 | 49 |
| Induced **Ecad** | | | 25 | 42 | | |
| Basal **Ecad** | | | | 47 | | |
| Induced **Lo.Ecad** | | | | | 40 | 49 |
| Basal **Lo.Ecad** | | | | | | 49 |

| | Induced **Ncad** | Basal **Ncad** | Induced **Pcad** | Basal **Pcad** | **C.Pcad** |
|---|---|---|---|---|---|
| Induced **Ncad** | 35 | 42 | 49 | 49 | 49 |
| Basal **Ncad** | | 47 | 49 | 49 | 49 |
| Induced **Pcad** | | | 35 | 42 | |
| Basal **Pcad** | | | | 47 | |
| **C.Pcad** | | | | | 43 |
